# A converged ubiquitin-proteasome pathway for the degradation of TOC and TOM tail-anchored receptors

**DOI:** 10.1101/2023.01.07.523072

**Authors:** Meijing Yang, Shuai Chen, Shey-Li Lim, Lang Yang, Jia Yi Zhong, Koon Chuen Chan, Zhizhu Zhao, Kam-Bo Wong, Junqi Wang, Boon Leong Lim

## Abstract

In plants, thousands of nucleus-encoded proteins translated in the cytosol are sorted to chloroplasts and mitochondria by binding to specific receptors of the TOC (translocon at the outer membranes of chloroplasts) and the TOM (translocon at the outer membranes of mitochondria) complexes for import into those organelles. The degradation pathways for these receptors are unclear. Here, we discovered a converged ubiquitin-proteasome pathway for the degradation of *Arabidopsis thaliana* TOC and TOM tail-anchored receptors. The receptors are ubiquitinated by E3 ligase(s) and pulled from the outer membranes by the AAA^+^ ATPase CDC48, after which a previously characterized cytosolic protein, TTOP, binds to the exposed transmembrane domains (TMDs) at the C termini of the receptors and CDC48, and delivers these complexes to the 26S proteasome.

## INTRODUCTION

The plant organelles chloroplasts and mitochondria are derived from two ancient endosymbiotic events, whereby a photosynthetic cyanobacterium and an aerobic prokaryote were separately engulfed by a eukaryotic cell. Over the intervening billion years of evolution, most genes from the endosymbiont genomes have been transferred to the host nucleus. Hence, most chloroplasts and mitochondrial proteins are encoded in the nucleus, synthesized in the cytosol, and imported into the organelles through the TOC (translocon at the outer membranes of chloroplasts) and the TOM (translocon at the outer membranes of mitochondria) complexes, respectively (Duncan et al. 2013; Shi and Theg 2013; Nakai 2018). Components of the TOC complex, including Toc33 and Toc159, are directed to the 26S proteasome by the ubiquitin-dependent chloroplast-associated protein degradation (CHLORAD) system (Ling et al. 2012; Ling et al. 2019). These TOC proteins are first ubiquitinated by SUPPRESSOR OF PPI1 LOCUS1 (SP1), an E3 RING ubiquitin ligase embedded in the chloroplast outer membrane (OM) (Ling et al. 2012). SP1 forms a complex with SP2, an Omp85-type β-barrel channel embedded in the OM, and CDC48A, a cytosolic AAA^+^ protein, at the chloroplast surface (Ling et al. 2019). Both SP2 and CDC48A are essential for the re-translocation of TOC components from the OM to the cytosol. SP2 is believed to play a conductance role, while CDC48A pulls the TOC proteins out of the membrane via ATP hydrolysis, after which ubiquitinated TOC proteins are targeted to the 26S proteasome for degradation (Ling et al. 2019). Yet, how ubiquitinated TOC proteins and the SP1/SP2/CDC48A complex are targeted to the 26S proteasome has been unclear. The CHLORAD system is critically important for chloroplast biogenesis, plant development, and stress responses (Ling et al. 2012; Ling and Jarvis 2015; Ling et al. 2019). Overexpression of *SP1* reduces the abundance of TOC complexes (Ling et al. 2012) and confers transgenic plants with higher resistance to salt, osmotic, and reactive oxygen species (ROS) stresses (Ling and Jarvis 2015). A reduction in the import of photosynthesis-related proteins can decrease the amount of ROS generated from chloroplasts during photosynthesis. Although SP1 is also present in mitochondria, whether it is involved in the ubiquitination of the TOM complex is unknown (Pan and Hu 2018). In addition, while the mitochondria-associated protein degradation (MAD) pathways of mitochondrial outer membrane (MOM) proteins are well characterized in yeast and mammalian cells (Zhang and Ye 2016; Zheng et al. 2019), the equivalent plant pathway(s) remain obscure. Here, we discovered a novel plant protein, which participates in ubiquitin-proteasome pathways for the degradation of *Arabidopsis thaliana* TOC and TOM tail-anchored receptors.

## RESULTS

### A novel cytosolic protein interacts with the C-terminal transmembrane domains of tail-anchored TOC and TOM receptors

We previously showed that Arabidopsis PURPLE ACID PHOSPHATASE2 (PAP2) plays a role in protein import into chloroplasts and mitochondria (Sun et al. 2012; Law et al. 2015; Zhang et al. 2016; Voon et al. 2021). Similar to Toc33 and Toc34 and to Tom20-2, Tom20-3, and Tom20-4, PAP2 is a tail-anchored (TA) protein anchored on the outer membranes of these two organelles via its hydrophobic C-terminal motif (Sun et al. 2012). PAP2 interacts with the precursor of the small subunit of Rubisco (pSSU) (Zhang et al. 2016) as well as the presequences of several MULTIPLE ORGANELLAR RNA EDITING FACTOR (pMORF) proteins (Law et al. 2015) and plays a role in their import into chloroplasts (Zhang et al. 2016) and mitochondria (Law et al. 2015), respectively. To further elucidate these processes, we first performed a yeast two-hybrid (Y2H) screen to identify PAP2-interacting proteins (Voon et al. 2021). One novel protein encoded by the gene At5g42220 drew our attention, as multiple clones were isolated during screening. The C-terminal transmembrane domain (TMD) of PAP2 was required for its interaction with At5g42220.1 (Fig. 1A). We named the protein encoded by At5g42220.1 <u>T</u>MD-binding protein for tail-anchored <u>o</u>uter membrane <u>p</u>roteins (TTOP). TTOP is an 879-amino-acid (aa) protein containing a ubiquitin-like (Ubl) domain at its N terminus (aa 24–95), while the rest of the protein lacks known conserved domains.

**Figure 1.**
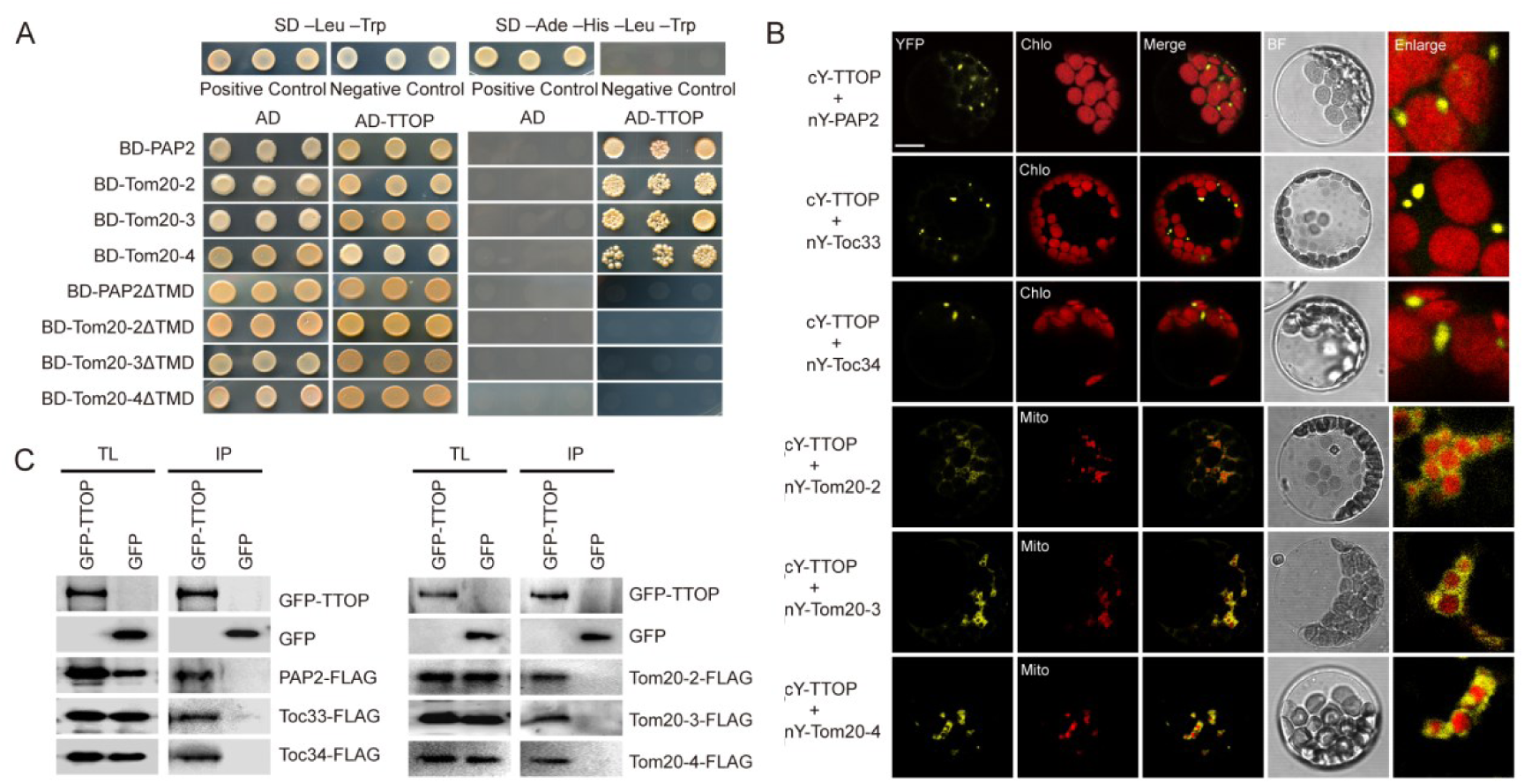
TTOP interacts with chloroplast and mitochondrial outer membrane tail-anchored proteins. **(A)** Yeast two-hybrid analysis of the interaction between TTOP and tail-anchored (TA) proteins. Yeast from the Y2HGold strain were co-transformed with pairs of the indicated constructs encoding TTOP and full-length or TMD-truncated (ΔTMD) TA proteins. The TMD-truncated TA proteins include PAP2ΔTMD (a.a. 1-614 & 637-656), Tom20-2ΔTMD (a.a. 1-182 & 201-210), Tom20-3ΔTMD (a.a. 1-174 & 193-202), and Tom20-4ΔTMD (a.a. 1-161 & 179-187). Co-transformation of pGADT7-T and pGBKT7-53 constructs, or pGADT7-T and pGBKT7-lam constructs, into yeast was employed as the positive and negative controls, respectively. **(B)** Bimolecular fluorescence complementation (BiFC) analysis of the interaction between TTOP and TA proteins. Protoplasts were transiently co-transfected with the indicated pairs of constructs, which encode the fusion proteins carrying complementary N- or C-terminal YFP fragments (nY or cY), respectively. Reconstitution of YFP fluorescence in protoplasts was detected by confocal microscopy; representative images are shown. Chlo, chloroplast autofluorescence. Mito, mitochondria marked with MitoTracker. BF, brightfield. Scale bar, 10 µm. **(C)** Co-immunoprecipitation (co-IP) of TA proteins in **(B)** by TTOP from protoplast extracts. Protoplasts were transiently co-transfected with constructs encoding FLAG-tagged TA proteins and GFP-TTOP or GFP. Protoplast total lysis (TL) and immunoprecipitates (IP) eluted from GFP-Trap agarose were subjected to immunoblotting analysis with anti-GFP and anti-FLAG antibodies, respectively.

Y2H analysis of the interaction of TTOP with other tail-anchored (TA) proteins on the outer membranes of chloroplasts and mitochondria revealed an interaction with Tom20-2, Tom20-3, and Tom20-4 (hereafter referred to together as Tom20-2/3/4) (Fig. 1A). The TMDs of the mitochondrial proteins were essential for their interactions with TTOP, as we observed no interaction between TTOP and TMD-truncated versions of Tom20-2/3/4 (Fig. 1A). We did not detect any interaction of TTOP with Toc33 and Toc34 (hereafter Toc33/34) by Y2H (Fig. S1), possibly because the BD-Toc33/34 fusion proteins could not enter the yeast nucleus, as the TA motifs from Toc33/34 and Tom20-2/3/4 have different properties (Sun et al. 2012). We then turned to bimolecular fluorescence complementation (BiFC) to confirm these interactions: TTOP fused to the C-terminal half of yellow fluorescent protein (cY) interacted with fusion proteins between the N-terminal half of YFP (nY) and Toc33/34, Tom20-2/3/4 and PAP2 (Fig. 1B), as determined by the reconstitution of YFP fluorescence. These interactions were dependent on their TMDs (Fig. S2), as there was no interaction between TTOP and TMD-truncated TA proteins (Fig. S3A). Notably, we did not detect interaction between the above TA proteins and TTOP fused to cY at its C terminus (Fig. S3B), indicating that a free TTOP C terminus is essential for these interactions. For TTOP interactions with Tom20-2/3/4, we occasionally observed YFP fluorescence surrounding mitochondria (Fig. 1B), which then appeared in the cytosol after a short duration (Fig. S4). These indicated that TTOP first interacted with Tom20-2/3/4 on the outer membrane of mitochondria and the interacting partners then migrated to the cytosol. For Toc33/34 and PAP2, we did not observe BiFC signals on the organellar outer membranes, but only in the cytosol (Fig. 1B). This observation was not due to a failure of targeting to the outer membranes, as GFP-TA fusions between the green fluorescent protein (GFP) and the TA proteins were successfully targeted to the organellar outer membrane (Fig. S5). These results suggested that TTOP interacts with cytosolic TA proteins carrying free TMDs but not with TA proteins whose TMDs are embedded in the outer membranes. In agreement with the BiFC results, co-immunoprecipitation (co-IP) assays from protein extracts of transiently co-transfected protoplasts confirmed the physical associations between TTOP and the above TA proteins (Fig. 1C).

### TTOP is essential for chloroplast biogenesis

To characterize the role of TTOP in organelle biogenesis and plant development, we first obtained two *ttop* mutants, SALK_128909 and SALK_151742, but these lines did not harbor a T-DNA insertion in *TTOP*, as determined by genotyping PCR. We therefore generated our own *TTOP* knock-out mutants using a highly efficient CRISPR/Cas9 system (Tsutsui and Higashiyama 2017), leading to the isolation of the *ttop-1* and *ttop-5* mutant alleles, with a single-base insertion of an A or T in the 43 bp of the first *TTOP* exon, respectively (Fig. S6A). Both alleles were predicted to introduce a premature stop codon 78 bp into the first exon. We selected Cas9-free *ttop-1* and *ttop-5* mutant lines and confirmed the absence of TTOP accumulation by immunoblotting with anti-TTOP antibodies (Fig. S6B). Both *ttop-1* and *ttop-5* mutants grew normally under standard growth conditions (Fig. S6C). We then attempted to generate transgenic lines overexpressing a construct encoding N-terminal GFP-tagged TTOP (GFP-TTOP), as adding a tag to the C terminus of TTOP affected its interaction with TA proteins in the BiFC assay. However, we failed to obtain lines that overexpressed *GFP-TTOP* when driven by either *UBIQUITIN10* (*UBQ10*) or the cauliflower mosaic virus *35S* promoter, suggesting that constitutive expression of *TTOP* might lead to lethality. Accordingly, we generated stable Arabidopsis transgenic lines expressing *GFP-TTOP* under the control of a dexamethasone (DEX)-inducible promoter (pTA7002*-GFP-TTOP*) (Aoyama and Chua 1997). These pTA7002*-GFP-TTOP* lines grew normally when sown on unadulterated MS medium but not on MS medium containing 10 μM DEX (Fig. 2A, B), confirming that constitutive overexpression of *GFP-TTOP* is early seedling-lethal, possibly due to an influence on chloroplast biogenesis, as indicated by the impairment of chloroplast development in cotyledons (Fig. 2C, D). When 5-day-old pTA7002*-GFP-TTOP* seedlings germinated on MS medium were transferred to MS medium containing 10 μM DEX for 3 days, their cotyledons turned yellow (Fig. 3A) and their chloroplasts degenerated, as evidenced by a decline in chloroplast and thylakoid size (Fig. 3B) observed during transmission electron microscopy (TEM) analysis. We confirmed the high accumulation of GFP-TTOP in pTA7002*-GFP-TTOP* lines by immunoblotting with anti-TTOP antibodies (Fig. 3B). Immunoblotting showed that higher TTOP protein accumulation reduced the abundance of outer envelope proteins in chloroplasts, without affecting the level of Tic40 (Fig. 3C). We failed to analyze mitochondrial proteins by immunoblotting, possible due to the low abundance of mitochondrial proteins in total protein extract and the lack of highly sensitive antibodies. We also tested the response of etiolated *ttop* seedlings to illumination, which induces chlorophyll biosynthesis and cotyledon opening in the wild type: Both *ttop* mutants also showed defects in cotyledon development and lower survival rates (Fig. 3D), as a consequence of impaired chloroplast biogenesis (Fig. 3E), upon exposure to light.

**Figure 2.**
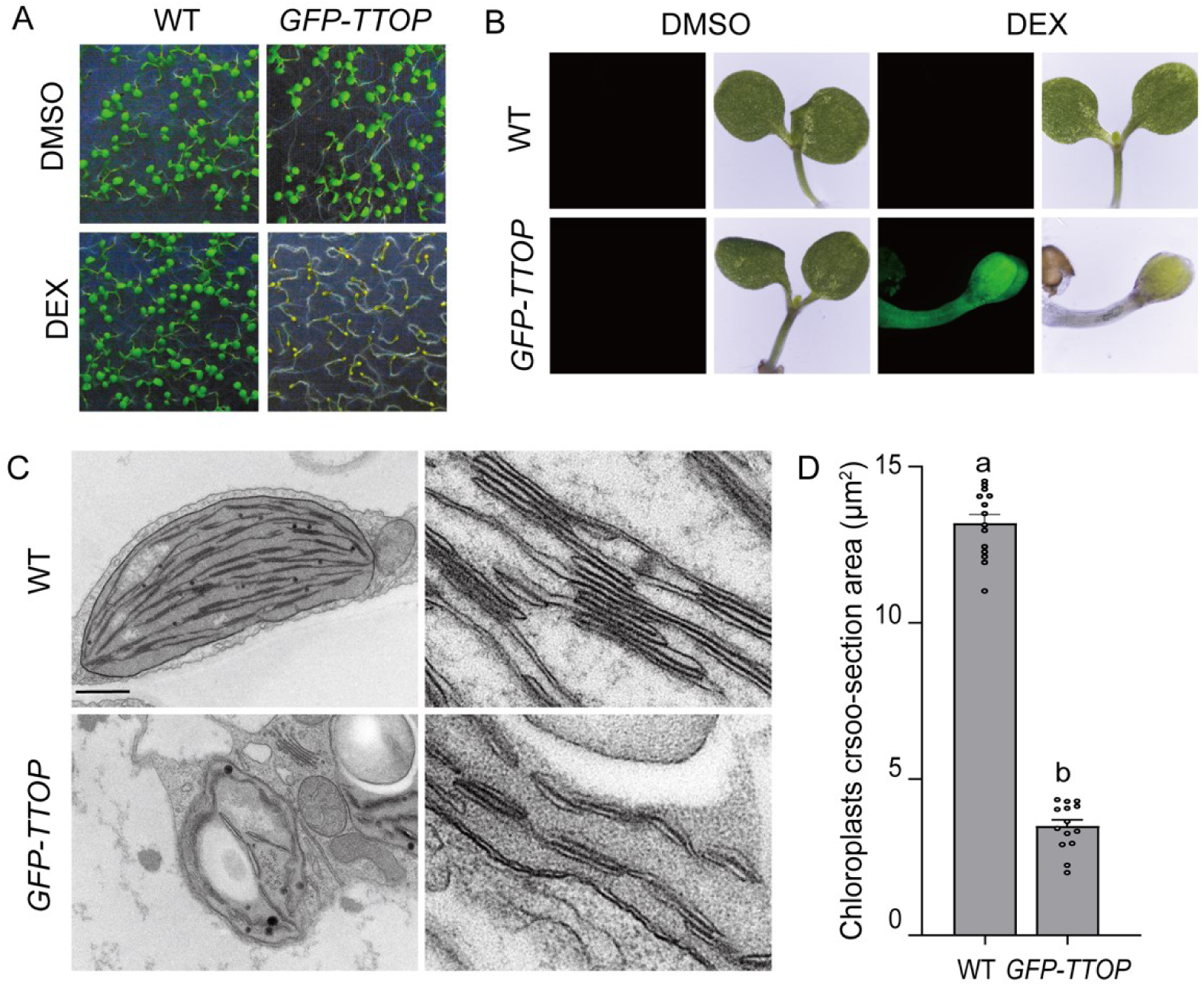
Overexpression of *TTOP* impairs chloroplast development. **(A and B)** Col-0 (WT) and pTA7002-*GFP-TTOP* seedlings were germinated on half-strength MS medium containing 10 µM DEX or DMSO (mock control treatment lacking the inducer) for 7 days. Typical seedling phenotypes are shown in **(A)**. GFP fluorescence in seedlings was detected by CLSM; typical images are shown **(B)**. **(C)** TEM analysis of the ultrastructure of cotyledon chloroplasts in the DEX-treated seedlings in **(A)**. Left, representative TEM micrographs of chloroplasts; right, thylakoid development. (**D**) Micrographs were used to estimate the cross-sectional area (bar chart) occupied by chloroplasts with ImageJ software. Scale bar, 1.0 µm. Bars show means ± SEM (n = 20 chloroplasts). Different letters indicate significant differences as analyzed by Tukey’s HSD test (P < 0.05).

**Figure 3.**
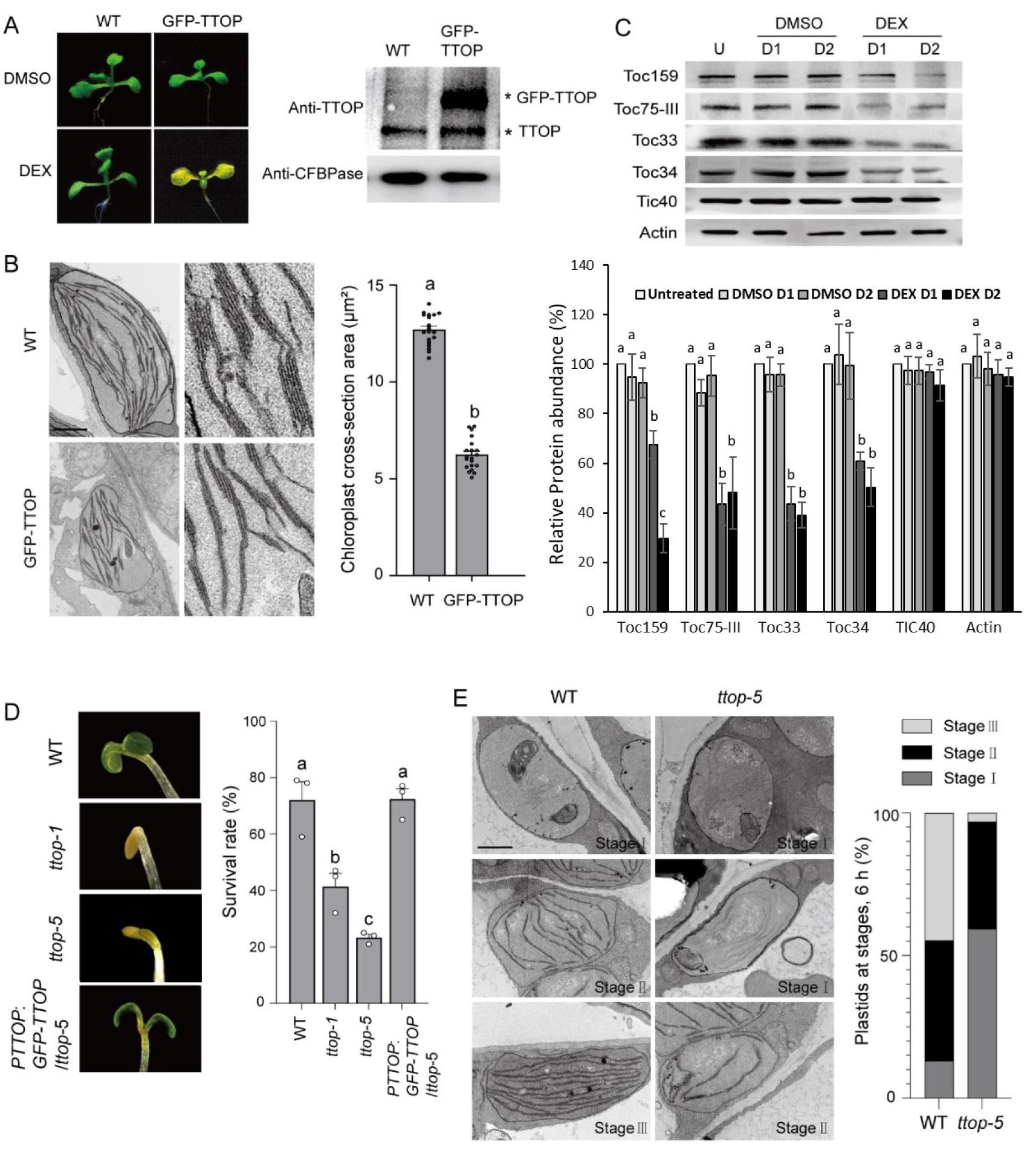
TTOP is essential for chloroplast biogenesis. **(A)** Left, representative images of 5-day-old pTA7002*-GFP-TTOP* and wild-type Col-0 (WT) seedlings grown on half-strength MS medium after 3 days of growth on half-strength MS medium containing 10 µM DEX or DMSO. Right, DEX-induced accumulation of GFP-TTOP as revealed by immunoblotting with anti-TTOP antibodies. Anti-cFBPase was used as a loading control. **(B)** Transmission electron microscopy (TEM) analysis of the ultrastructure of chloroplasts in cotyledons of seedlings shown in **(A)**. Micrographs show chloroplasts (left) and thylakoid development (right). Scale bar, 1.0 µm. Bar chart shows estimates of chloroplast cross-sectional area determined from the micrographs using ImageJ software. **(C)** Immunoblotting analysis of TOC receptors in total leaf proteins from pTA7002*-GFP-TTOP* plants subjected to DEX or DMSO treatment for 1 (D1) or 2 days (D2), or untreated (U). Anti-actin was used as a loading control. Untreated samples = 100%. Band of untreated samples were normalized **(D)** Results of de-etiolation of *ttop-1*, *ttop-5*, *ttop-5 pTTOP:GFP-TTOP*, and WT seedlings, grown in the dark for 6 days, upon transfer into continuous light. After 2 days of illumination, cotyledon phenotypes (left) and plant survival rates (right) were recorded. **(E)** Left, TEM analysis of the ultrastructure of cotyledon plastids in different genotypes after 0, 6, and 24 hours of illumination as in **(D)**. Right, proportions of plastids at each of three developmental stages estimated after 6 hours of illumination. Scale bar, 1.0 µm. Bars show means ± SEM (*n* = 20 chloroplasts in B, *n* = 3 experiments in C, *n* ≥ 30 images in D, E). Different letters indicate significant differences as analyzed by Tukey’s HSD test (*P* < 0.05).

### TTOP is expressed in actively dividing tissues

To examine the expression pattern of *TTOP*, we generated transgenic lines harboring a transgene driving the expression of the *β-GLUCURONIDASE* (*GUS*) reporter gene under the control of the *TTOP* promoter (*pTTOP*:*GUS*) (Fig. 4A-I). GUS staining was strong in root and shoot meristems (Fig. 4A–D), which are characterized by actively dividing cells with high energy demand. The *TTOP* promoter was active in young leaves but not in mature leaves (Fig. 4E) and was also active in trichomes, flowers, and pollen (Fig. 4F–I). To determine the cellular localization of TTOP, we generated *ttop-5 pTTOP*:*GFP-TTOP* transgenic plants by transforming the construct *pTTOP*:*GFP-TTOP*, driving *TTOP* expression from the *TTOP* promoter, into *ttop-5* plants. The transgene complemented the *ttop-5* mutant, as evidenced by the rescue of the cotyledon phenotypes (Fig. 3D) upon de-etiolation. These results hinted that *TTOP* expression is strictly controlled. Confocal laser scanning microscopy (CLSM) analysis of stable Arabidopsis *ttop-5 pTTOP*:*GFP-TTOP* seedlings showed that GFP-TTOP accumulated in the cytosol of all tissues tested, including mesophyll cells, the epidermis, the hypocotyl, and the roots, and was highly abundant in the root and shoot meristems and the first true leaves of 6-day-old seedlings (Fig. 4J). The cytosolic location of TTOP is also confirmed in protoplasts (Fig. S5).

**Figure 4.**
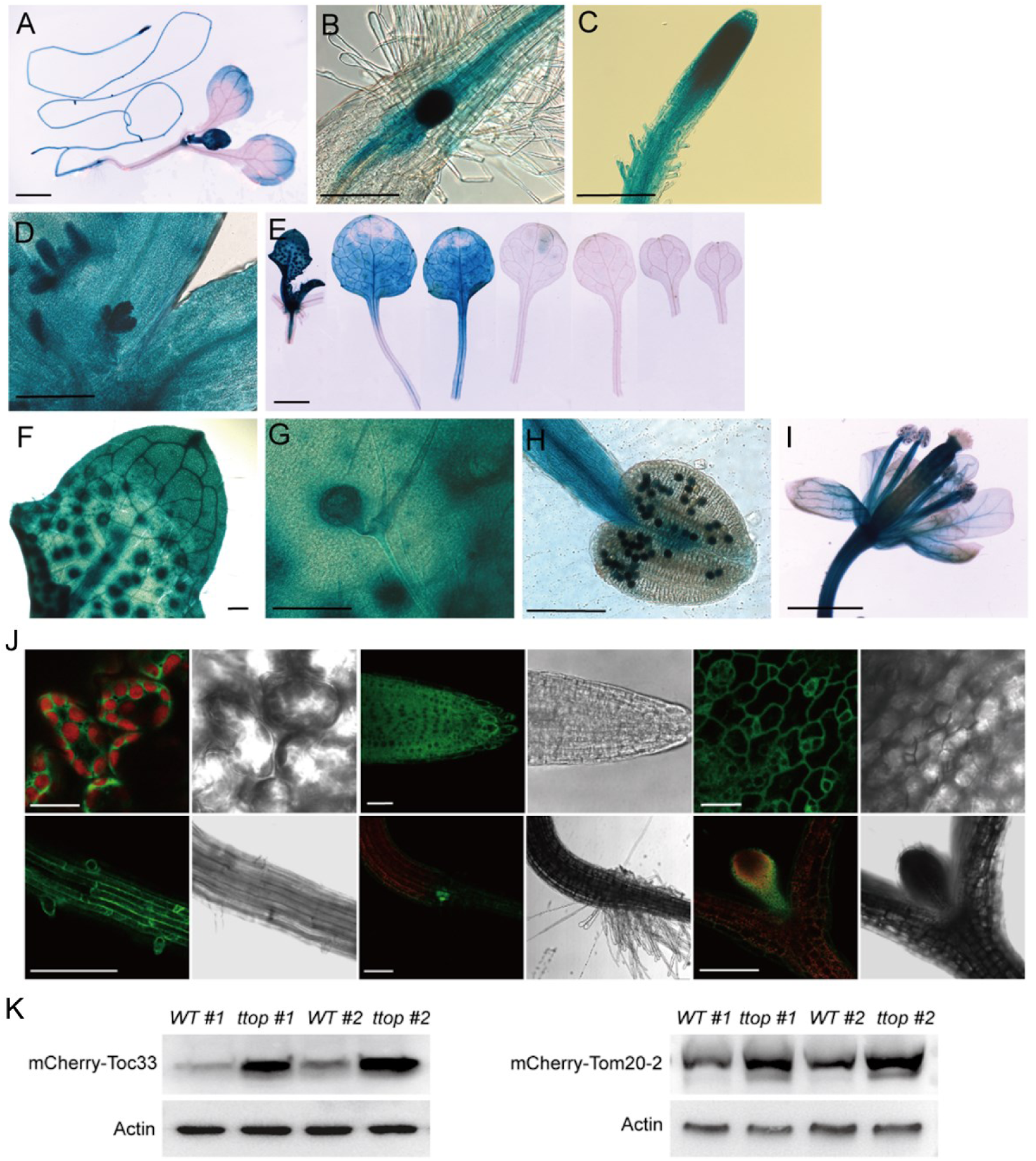
*In planta* expression patterns of TTOP. Expression patterns of *TTOP* at different developmental stages in *pTTOP:GUS* plants. Ten-day-old seedlings **(A-D, F, G)** and 23-day-old plants **(E)** from *pTTOP:GUS* transgenic lines were used for GUS staining. Decreases in *TTOP* expression were observed in older tissues, particularly in leaves **(A, E)**, with high *TTOP* expression in newly grown leaves and no *TTOP* expression in fully grown leaves **(E;** left to right, youngest to oldest leaves). High *TTOP* expression was also observed in meristematic regions, including the root-hypocotyl junction **(B)**, root meristem **(C)**, and shoot meristem **(D)**, and in other tissues such as trichome bases **(F** and **G)**, mature pollen **(H)**, and flowers **(I).** Scale bar in A, E, I, 2.0 mm. Scale bar in B, C, D, F, G, H, 200 µm. (**J)** Tissue expression pattern and subcellular localization of TTOP in 6-day-old *ttop-5 pTTOP:GFP-TTOP* transgenic seedlings. TTOP accumulates in mesophyll cells, roots, and epidermis. TTOP has a cytosolic localization in different tissues. Red signal is chlorophyll autofluorescence. Brightfield images are shown for reference. Scale bar in the upper panels, 20 µm. Scale bar in the below panels, 200 µm. **(K)** Transgenic *ttop-5* mutant plants expressing mCherry-Toc33 or mCherry-Tom20-2 were crossed with WT plants to obtain WT plants expressing mCherry-Toc33 or mCherry-Tom20-2 for comparisons of mCherry-Toc33 or mCherry-Tom20-2 abundance between *ttop-5* and WT. Two individual homozygous lines for each *ttop* and WT genotype were tested by immunoblotting with anti-mCherry antibodies, which showed more mCherry-Toc33 and mCherry-Tom20-2 in the *ttop-5* mutant than the WT. Anti-actin was used as a loading control.

### TTOP participates in the CHLORAD pathway

As TTOP interacted with TA proteins targeted to chloroplasts and mitochondria, we generated *p35S:mCherry-Toc33* and *p35S:mCherry-Tom20-2* transgenic plants in the *ttop-5* mutant background. We then crossed these transgenic lines with wild-type (WT) plants and selected homozygous *p35S:mCherry-Toc33* and *p35S:mCherry-Tom20-2* transgenic plants in the WT background from their progeny. With this crossing strategy, we thus obtained *p35S:mCherry-Toc33* and *p35S:mCherry-Tom20-2* transgenic plants in the WT and *ttop-5* backgrounds carrying the same T-DNA insertion events, allowing us to compare mCherry-Toc33 and mCherry-Tom20-2 protein abundance. The abundance of mCherry-Toc33 and mCherry-Tom20-2 was the same in the two WT lines (Fig. 4K). Importantly, mCherry-Toc33 and mCherry-Tom20-2 abundance was much higher in *ttop-5* relative to the WT lines, as evidenced by immunoblotting with anti-mCherry antibodies (Fig. 4K). We thus concluded that TTOP may contribute to the degradation of Toc33 and Tom20-2, which prompted us to explore whether TTOP participates in the CHLORAD pathway. BiFC analysis showed that TTOP interacts with SP1, CDC48A, and the 26S proteasome subunits REGULATORY PARTICLE NON-ATPASE6 (RPN6), RPN10, RPN12, and RPN13a (Fig. 5A), but not with SP2. In agreement with this, co-IP assays from transiently transfected protoplast extracts showed that TTOP can pull down SP1, CDC48, RPN6, RPN10, RPN12 and RPN13 (Fig. 5B). Co-transfection of *mCherry-TTOP*, *cY-CDC48A*, and *nY-Toc33/Toc34* constructs in Arabidopsis protoplasts resulted in the reconstitution of YFP fluorescence and co-localization of YFP with mCherry fluorescence in the cytosol, indicating that these proteins co-localize (Fig. 6). Likewise, co-transfection of *cY-CDC48A*, and *nY-Tom20-2/Tom20-3/Tom20-4* constructs in protoplasts also demonstrated their co-localization in the cytosol and exhibited a network appearance (Fig. 7), which disappeared when TTOP was co-expressed (Fig. 6). CDC48A interacted with Toc33/34 and Tom20-2/3/4 at the periphery of chloroplasts and mitochondria, as indicated by BiFC assays (Fig. 7). We repeated the BiFC assays in protoplasts prepared from *ttop-5* plants and observed the same results (Fig. S7), indicating that TTOP is not essential for the interaction of CDC48A with Toc33/34 or Tom20-2/3/4.

**Figure 5.**
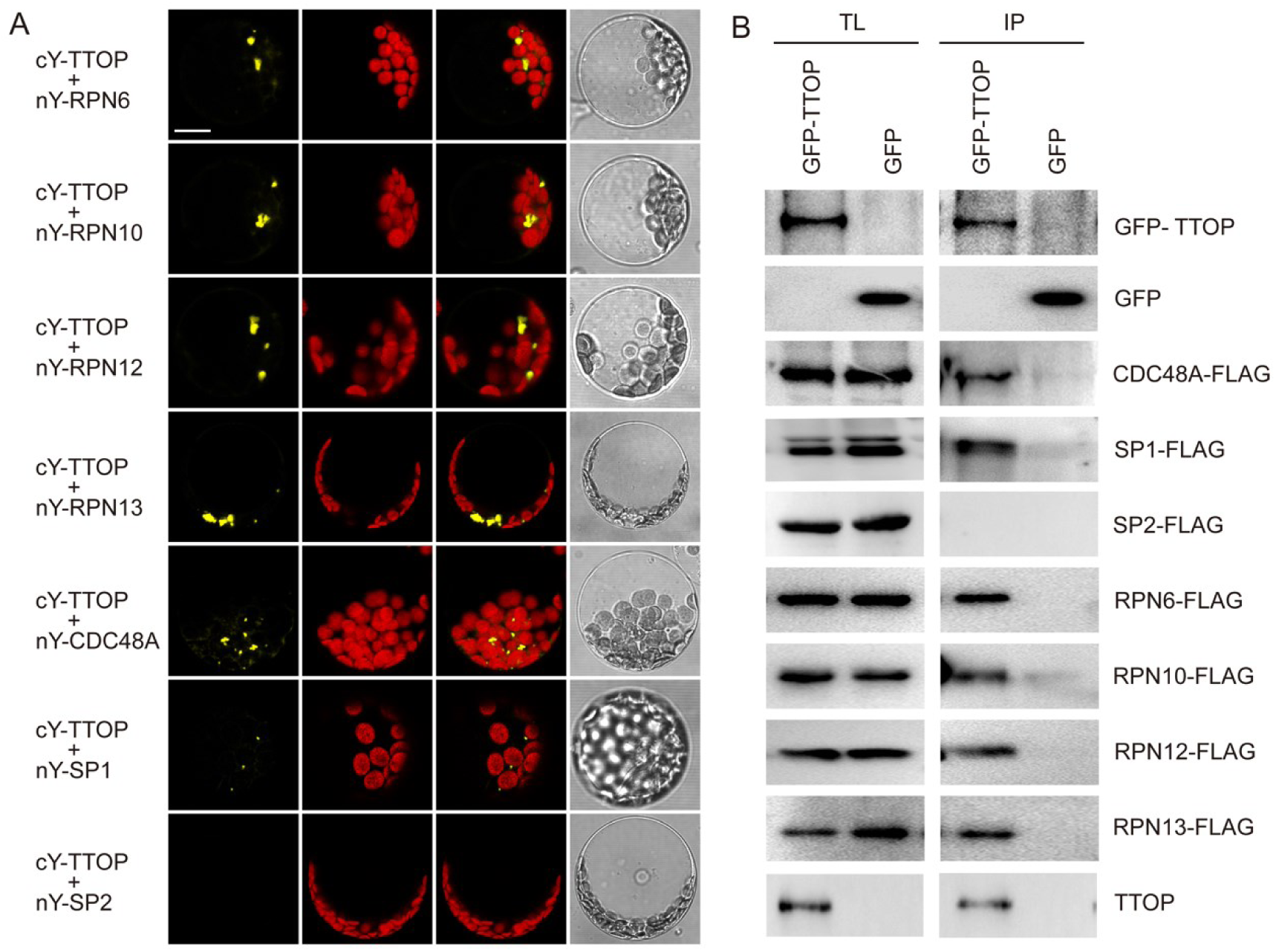
TTOP, chloroplast, and mitochondrial TA proteins, CDC48A, and 26S proteasome subunits form complexes *in vivo*. **(A)** BiFC analysis of the interactions between TTOP and CDC48A, SP1, SP2, and 26S proteasome subunits. Protoplasts were co-transfected with the indicated pairs of constructs encoding fusion proteins with cY or nY. Reconstitution of YFP fluorescence in protoplasts was detected by confocal microscopy. Chlo, chloroplast autofluorescence. BF, brightfield. Scale bar, 10 µm. **(B)** Co-IP of CDC48A, SP1, or 26S subunits with TTOP from protoplast extracts. Protoplasts were co-transfected with constructs encoding GFP-TTOP or GFP and constructs encoding FLAG-tagged CDC48A, SP1, or 26S proteasome subunits. Protoplast extracts and elution proteins from GFP-Trap agarose were subjected to immunoblotting analysis with anti-GFP and anti-FLAG antibodies as described above.

**Figure 6.**
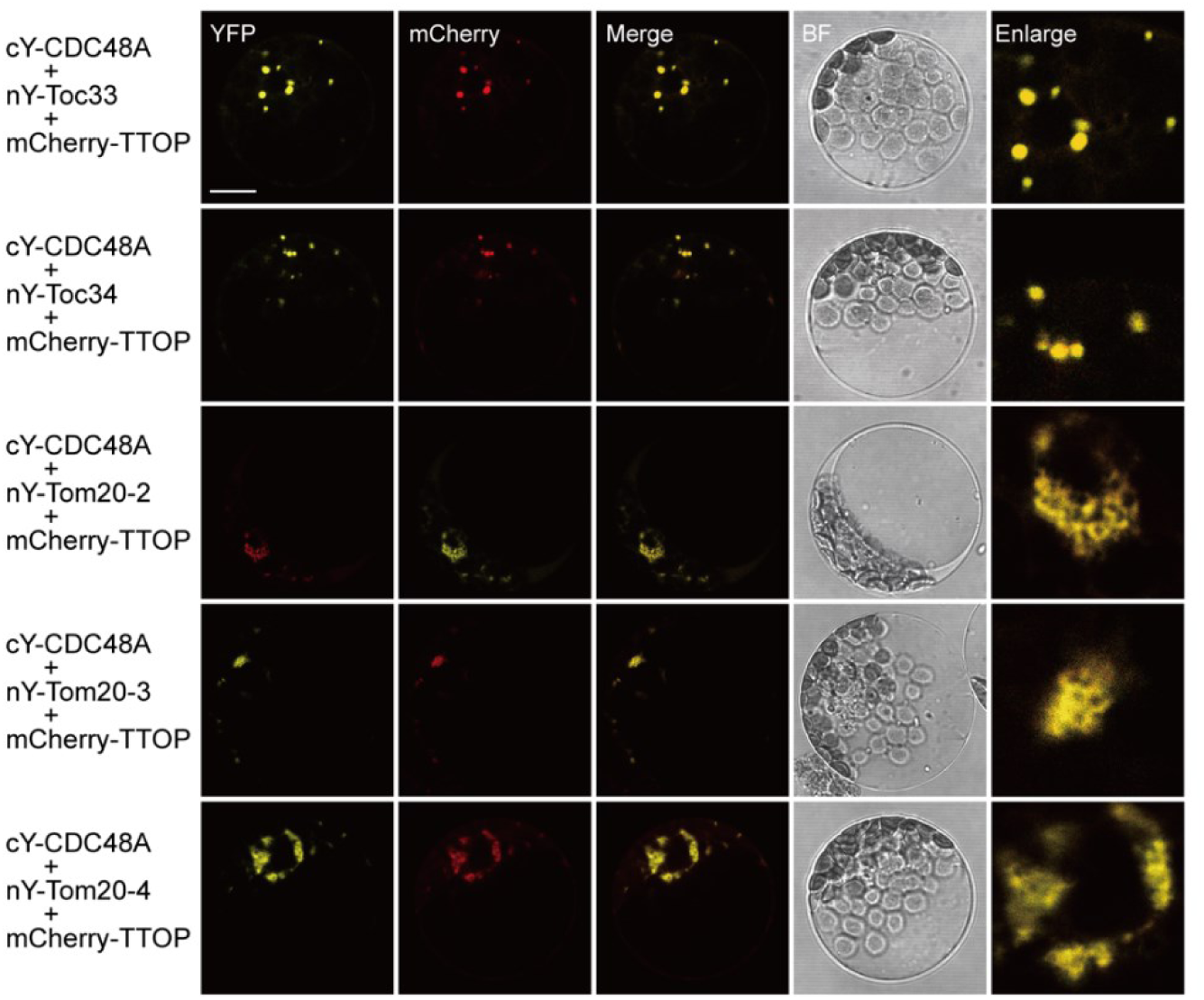
Co-localization of TTOP, CDC48A, and chloroplast or mitochondrial outer membrane TA receptors in protoplasts. Protoplasts were co-transfected with constructs encoding TTOP fused to mCherry at its N terminus or encoding CDC48A and TA proteins fused to nY or cY at their N termini. YFP and mCherry fluorescence in protoplasts was analyzed by CLSM; representative confocal images are shown. Overlap of YFP and mCherry signals is shown in protoplasts, indicating the co-localization of TTOP and CDC48A with TA proteins. BF, brightfield. Scale bar, 10 µm.

**Figure 7.**
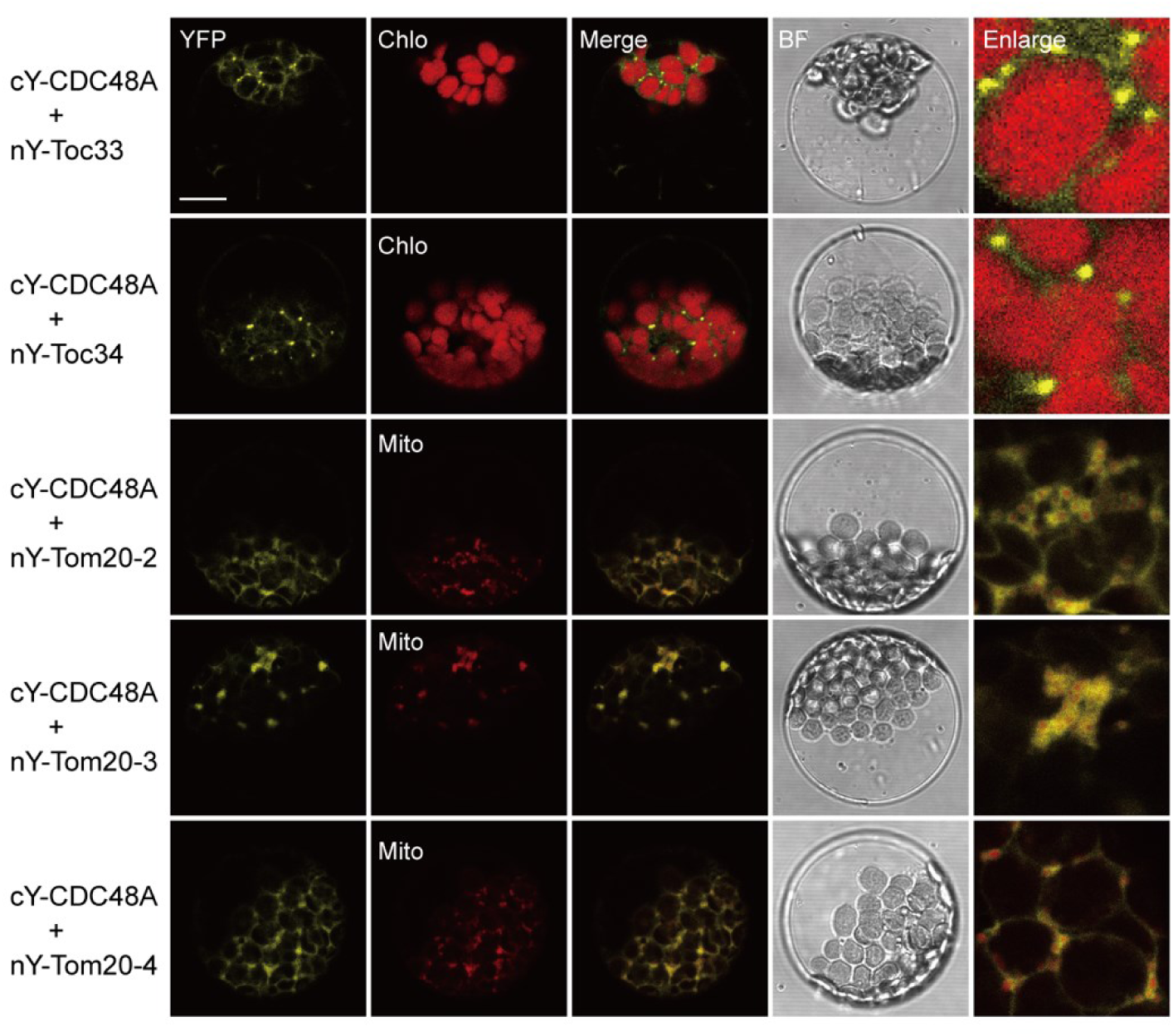
BiFC analysis of the interaction between CDC48A and the TA receptor proteins. Protoplasts were co-transfected with the indicated pairs of constructs encoding CDC48A or TA proteins carrying nY or cY in their respective N termini. Reconstitution of YFP fluorescence in protoplasts was detected by CLSM, and the representative confocal images are shown. Chlo, chloroplast auto-fluorescence; Mito, mitochondria marked with MitoTracker. BF, brightfield. Scale bar, 10 µm.

### TTOP may participate in a ubiquitin-dependent MAD pathway

Next, we tested whether SP1 can ubiquitinate Toc33 and Tom20-3 by *in vitro* ubiquitination assays. Recombinant Arabidopsis UBIQUITIN-ACTIVATING ENZYME1 (UBA1), UBIQUITIN CONJUGATING ENZYME8 (UBC8), and ubiquitin (Fig. S8), recombinant SP1flex, TTOP (Fig. S9) and Toc33 and Tom20-3 (Fig. S10) were purified in *E. coli*. GST-SP1RING, but not SP1flex, self-ubiquitinated (Fig. S11A). These data showed that the SP1RING domain is able to ubiquitinate its GST fusion partner, whereas SP1flex which contains SP1RING domain cannot self-ubiquitinated itself. SP1flex also ubiquitinated full length Toc 33 (Toc33FL, a.a. 1-297) and Toc33 without the C-terminus (Toc33NC, a.a. 1-251) (Fig. S11B). GST-TTOP, but not GST alone, pulled down both non-ubiquitinated (Fig. S11C) and ubiquitinated Toc33FL (Fig. S11D), but not Toc33NC in either form (Fig. S11E), indicating that SP1 ubiquitinates the receptor domain of Toc33, whereas TTOP binds to the TMD motif of Toc33, independent of its ubiquitination status. SP1flex/UBC8 did not ubiquitinate Tom20-3 without the C-terminus (Tom20-3NC, a.a. 1-174) or TTOP (Fig. S11F). Unidentified E2/E3 ligase(s) in seedling extracts appeared to ubiquitinate His-Tom20-3NC, which was not pulled down by GST-TTOP due to the lack of the TMD motif of Tom20-3 (Fig. S11G). Although we could not test the ubiquitination of full length Tom20-3 as we failed to express it in *E. coli*, the TMD motifs of Tom20-2/3/4 cannot be ubiquitinated as they do not contain any lysine residues. Therefore, the ubiquination sites must be located at the cytosolic domain of Tom20-3. Unknown E3 ligase(s) in the plant extracts may therefore ubiquitinate Tom20-3, or SP1 at the mitochondrial OM (Pan and Hu 2018) may still act as the cognate E3 ligase of Tom20-3, as our *in vitro* experiments did not test all 37 Arabidopsis E2 ligases. Further studies are required to delineate the details of the ubiquitin-dependent MAD pathway.

### TTOP binds to RPN13 of the 26S Proteosome via its N-terminal UBL domain

Using the TTOP protein sequence to PBLAST the proteins in the human genome, the protein with the highest protein sequence identity was human BAG6 (Chio et al. 2017; Shao et al. 2017). The length of BAG6 (1132 a.a.) is much longer than TTOP (879 a.a.) and they only share 23% protein sequence identity, mainly at their N-terminal UBL domains and their C-terminal BAG domain (Mock et al. 2015). In mammalian system, the Bag6/Ubl4A/Trc35 complex directs mislocated polypeptides on the ER membrane toward 26S proteasome for degradation to avoid protein aggregations in the cytosol (Hessa et al. 2011; Wang et al. 2011). Besides BAG6, many UBL-containing proteins are involved in shuttling ubiquitinated substrates to 26S proteasomes and some UBL domains were shown to bind to RPN subunits (Chen et al. 2016; Shi et al. 2016). Here, we employed AlphaFold2 (Jumper et al. 2021; Mirdita et al. 2021) to predict the interaction between the UBL domain of TTOP with the RPN proteins we identified by BiFC. As predicted by AlphaFold2, the UBL domain of TTOP could interact with the pleckstrin-like receptor for ubiquitin (PRU) domain of RPN13 (Fig. S12D and Fig. 8A). The interaction between these two domains was confirmed by ITC experiment and the affinity was determined to be 29 μM (Fig. 8D). In addition, sequence alignment with the UBL domain of BAG6 showed that TTOP UBL contains the RNF126_NZF interacting residues (Krysztofinska et al. 2016), suggesting that TTOP UBL may have a zinc finger (ZF) binding ability (Fig. S13A). SP1 RING is a ZF protein with the E3 ligase function (Pan and Hu 2018; Ling et al. 2019) and Alphafold2 predicted that the UBL domain of TTOP could bind to SP1 RING, too (Fig. S13B). By contrast, the C-terminus of TTOP contains a BAG domain that contains conserved Ubl4A interaction residues (Mock et al. 2015) and this domain was predicted to be able to bind human Ubl4A by AlphaFold2 (Fig. S14).

**Figure 8.**
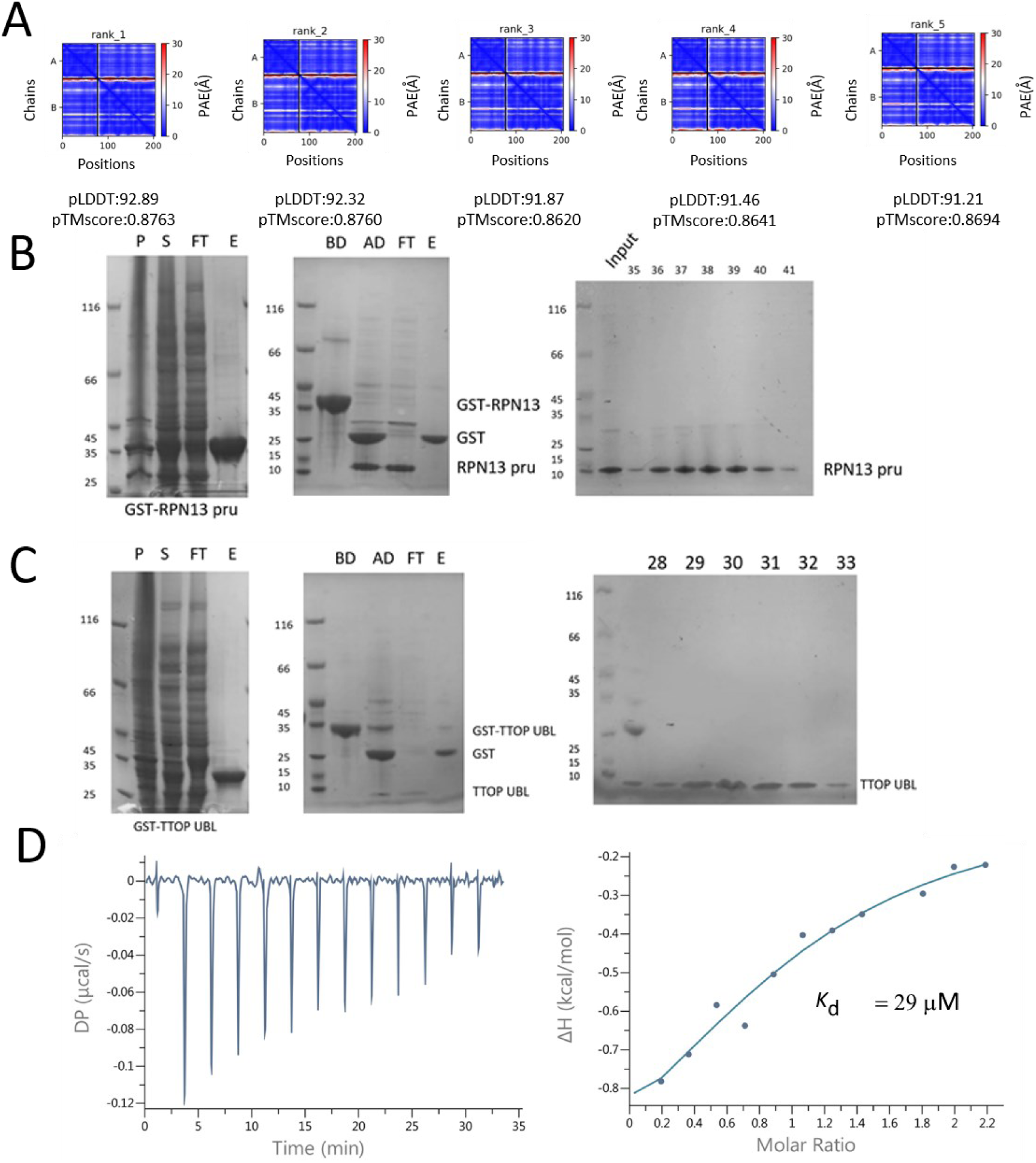
TTOP UBL domain could interact with RPN13 pru domain. By AlphaFold2 prediction, the high value of pLDDT and pTMscore and the low value of PAE showed that TTOP UBL could interact with the pru domain of RPN13. Recombinant TTOP UBL domain (B) and RPN13 pru domain (C) were purified by using GST fusion protein purification, on-column digestion and HiLoad 26/200 Superdex 200pg column. Isothermal titration calorimetry (ITC)-based measurement of the binding affinity between the UBL domain of TTOP and the pru domain of RPN13 at a 1:1 ratio is determined to be 29µM (D).

### Heat treatment induces TTOP-dependent clearance of mCherry-Toc and mCherry-Tom20-2 in planta

We noticed that heat treatment can induce patch formation of GFP-TTOP in protoplasts (Fig. S15A). Then we generated double expression lines by transforming the homozygous DEX-GFP-TTOP line with the *p35S:mCherry-Toc33* and *p35S:mCherry-Tom20-2* constructs to carry out heat-induction experiment. These lines expressed mCherry-Toc33 and mCherry-Tom20-2 constitutively at high levels. GFP-TTOP was induced by 24h Dex treatment in some 8-day-old seedlings and then subject to 30 min heat treatment at 45°C. While the mCherry signals could still be seen immediately after the heat treatment, the signals of mCherry-Toc33 and mCherry-Tom20-2 gradually disappeared (1h and 3h post treatment) when GFP-TTOP was present, but remained when GFP-TTOP was absence (Fig. 9 and Fig. S16). These in planta data showed that GFP-TTOP is required for the degradation of mCherry-Toc33 and mCherry-Tom20-2. Heat treatment was also shown to induce mRNA expression of TTOP (Fig. S15B) and the *ttop-5* mutants were less heat-resistant than the WT seedlings (Fig. S15C).

**Figure 9.**
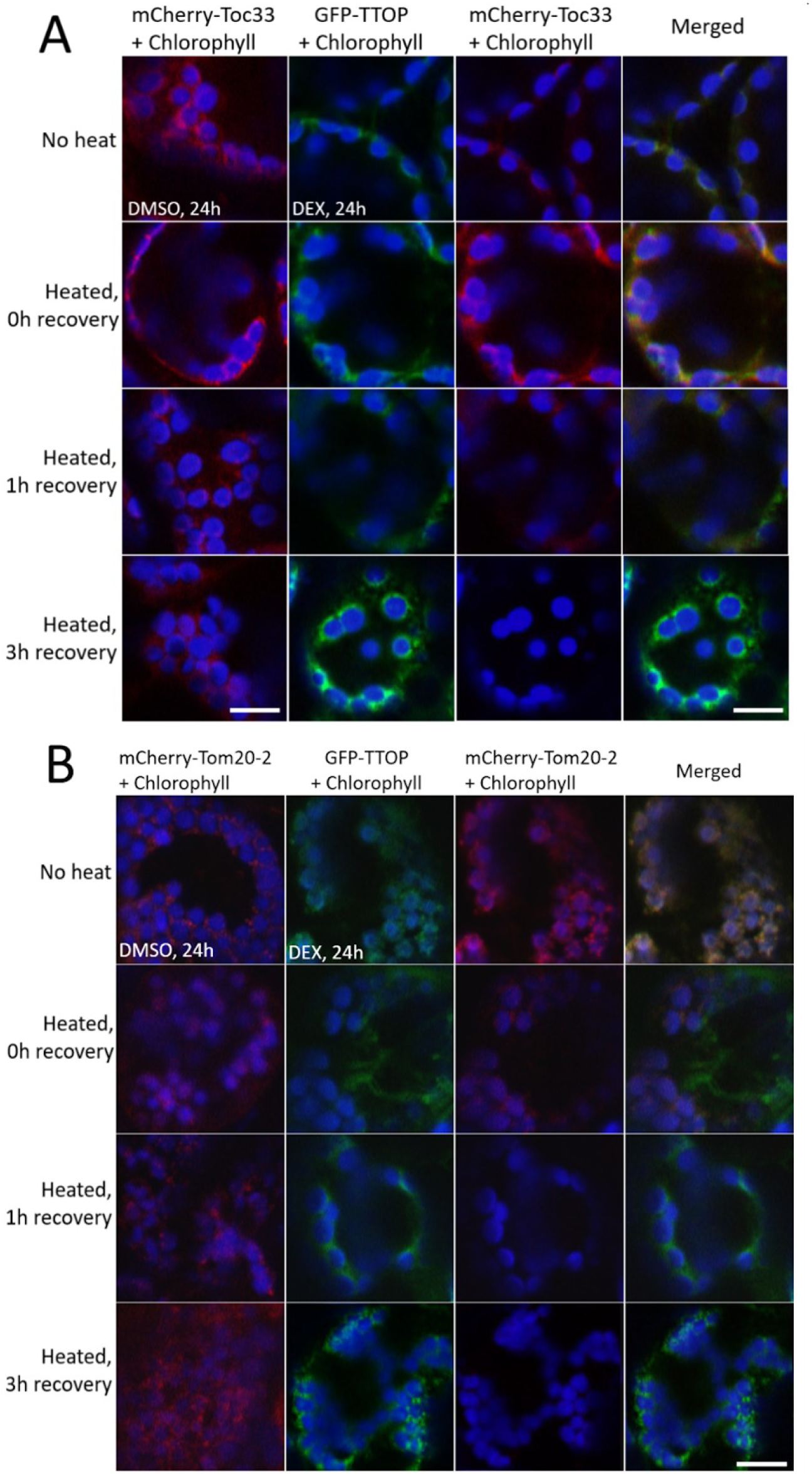
TTOP assists the degradation of Toc33 and Tom20-2 in planta. The transgenic DEX-GFP-TTOP line was transformed with 35S: mCherry-Toc33 (A), and 35S: mCherry-Tom20-2 (B), respectively. 25μM DEX were sprayed to the seedlings at 8-day-old. Some seedlings were treated with DMSO as negative controls. After 24 h, seedlings were placed in 45°C incubator for 30 minutes (heated) or kept in R.T. (no heat). Heat-treated seedlings were allowed to recover in R.T. for 1 and 3 hours prior to the imaging. Green: GFP-PFP; Red: mCherry-Toc33; Blue: Chlorophyll auto-fluorescence. Bars = 20 μm.

## DISCUSSION

In this study, we discovered a converged pathway for the delivery of ubiquitinated tail-anchored receptors of the TOC and TOM complexes to the 26S proteasome for degradation (Fig. 10). In CHLORAD, a system for the clearance of the TOC complex by the ubiquitin-proteasome system, the E3 ligase SP1 mediates the ubiquitination of the TOC complex, while SP2 and the AAA^+^ ATPase CDC48A act as a conduit and a motor, respectively, to retrotranslocate the ubiquitinated TOC complex and SP1 out of the chloroplast OM (Ling et al. 2019). However, how the retrotranslocated, ubiquitinated TOC proteins are delivered to the 26S proteasome was not clear. Here, we showed that TTOP is the cytosolic shuttling factor responsible for the delivery of the ubiquitinated TOC proteins to the 26S proteasome. Our observations fit with the previous findings that CDC48 plays a role in the retrotranslocation of ubiquitinated Toc33 and SP1 from chloroplasts to cytosol (Ling et al. 2019). TTOP interacts with TOC proteins and SP1 in the cytosol after they are retrotranslocated from the chloroplast OM by CDC48. In addition, our data suggested the existence of a similar MAD pathway for the removal of Tom20 from the mitochondrial OM, which involves uncharacterized E2/E3 ligases, CDC48A, and TTOP. In this pathway, ubiquitinated Tom20 receptors of the TOM complexes are retrotranslocated from the OM to the cytosol, followed by the delivery of the complex to the 26S proteasome by TTOP. Ubiquitination of the TOC and TOM receptors is not required for TTOP binding; hence, the binding takes place after these receptors are retrotranslocated from the OMs to the cytosol. During TOC biogenesis, cytosolic Arabidopsis ankyrin repeat protein 2A (AKR2A) binds to the TMDs of newly synthesized Toc33 and Toc34, but not that of Tom20-2, when they are released from ribosomes to maintain the nascent TOC receptors from forming aggregates or binding to non-specific hydrophobic proteins before their insertion into the chloroplast OM (Bae et al. 2008; Kim et al. 2019). Our BiFC data showed that TTOP first interacted with Tom20-2/3/4 on the OM of mitochondria (Fig. 1B), and the complexes migrated to cytosol after a short duration (Fig. S4). Future research is required to identify the E2/E3 ligases that ubiquitinate Tom20 receptors and the role of CDC48 in their retrotranslocation. For Toc33/34, we did not observe BiFC signals on the chloroplast outer membranes, but only in the cytosol (Fig. 1B). Perhaps the retrotranslocation of Toc33/34 happened more efficiently than that of Tom20-2/3/4 in the BiFC experiments. Nonetheless, while both AKR2A and TTOP can bind to the TMDs of Toc33 and Toc34, they might bind to the receptors at different processes: the biogenesis and degradation stages of the TOC complexes, respectively.

**Figure 10.**
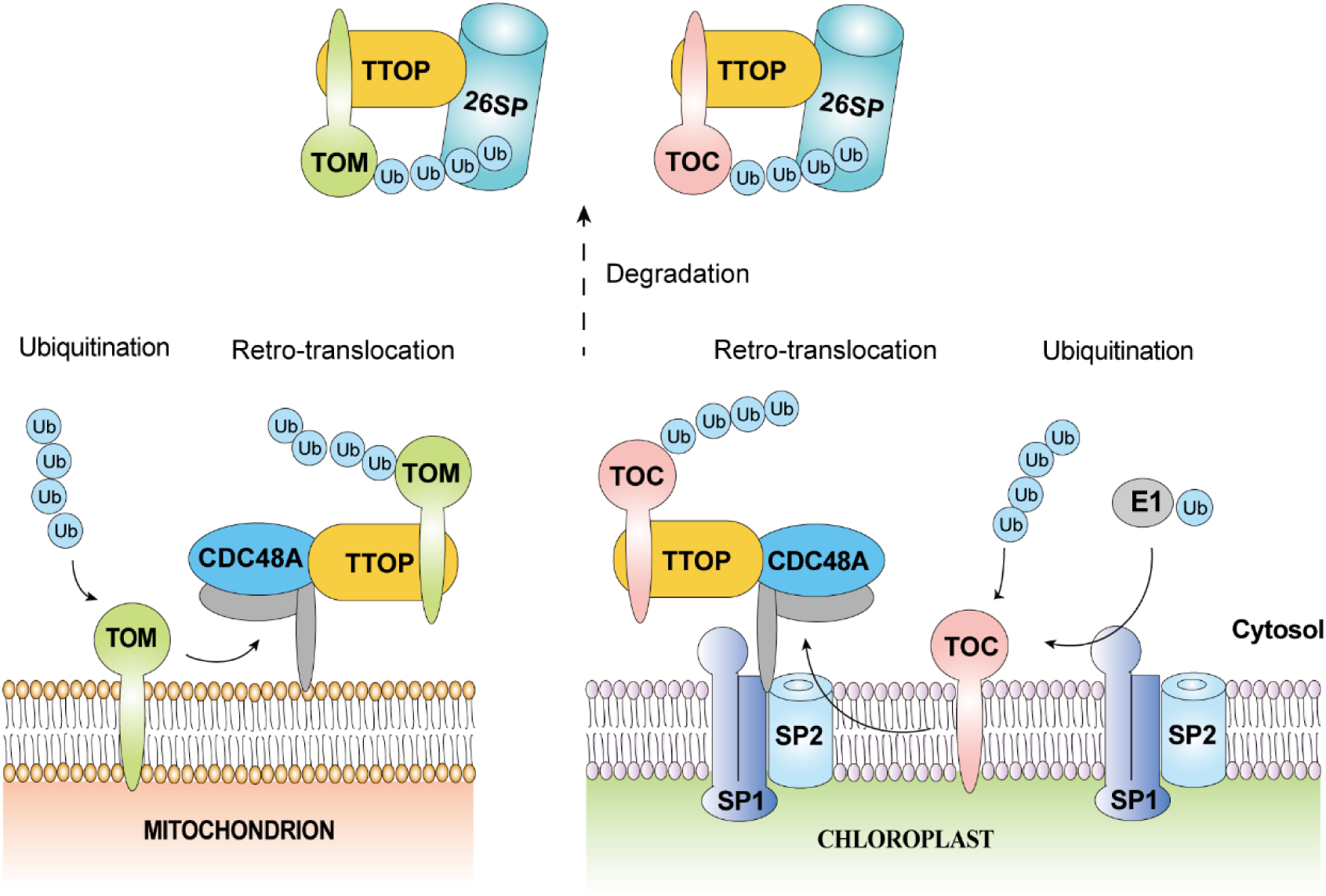
Proposed model for the role of TTOP in a converged pathway for the degradation of tail-anchored receptors of TOC and TOM translocases. In the CHLORAD proteolytic system, TOC components of the chloroplast protein import machinery are selectively removed by the SP1-SP2-CDC48A complex. After the TOC tail-anchored receptors are ubiquitinated by SP1, SP2 and CDC48A act as a conduit and a molecular motor, respectively, to mediate retrotranslocation of TOC complex out of the membrane. Similarly, tail-anchored receptors of TOM complexes are also ubiquitinated by an unknown E3 ubiquitin ligase and presumably pulled out from the outer envelope membrane (OEM) by CDC48A. Cytosolic TTOP then binds to the transmembrane domains (TMDs) of the retrotranslocated TOC/TOM receptors, masking the hydrophobic aggregation-prone TMDs and stabilizing ubquitinated TOC/TOM complexes in the cytosol. TTOP, via its interaction with RPN subunits of the 26S proteasome, then delivers the ubiquitinated receptors to the 26S proteasome for degradation. Association of TTOP with both CDC48A and RPN subunits of the 26S proteasome might allow efficient delivery of ubiquitinated receptors to the proteasome, thereby avoiding the risk of protein aggregation and enhancing degradation efficiency.

Besides the UBL domain at its N-terminus and the BAG domain at its C-terminus, the other region of TTOP does not show any sequence homology to any known structures. In the mammalian Bag6/Ubl4A/Trc35 complex, the C-terminal BAG domain of BAG6 interacts with UblA4 (Mock et al. 2015), which in turn interacts with the small glutamine-rich tetratricopeptide repeat-containing protein alpha (SGTA), and the latter binds TMD directly (Shao et al. 2017; Guna and Hegde 2018). Different from the BAG6 in the mammalian system, TTOP can bind to TMDs of the TOC/TOM TA receptors directly. Although the BAG domain at the C-terminus of TTOP was predicted to be able to interact with human UblA4 by Alphafold2 (Fig. S14), PBLAST search could not find any homolog of UblA4 in Arabidopsis and the closet homolog of SGTA are proteins containing the tetratricopeptide repeat (TPR) domains, including HOP1(Fellerer et al. 2011), HOP2, HOP3 (Fernández-Bautista et al. 2017), OM64 (Panigrahi et al. 2014) and Toc64 (Sommer et al. 2013), of which they only share homology with SGTA at their TPR domains. It might be interesting to investigate whether these proteins could bind to TMDs and play a role similar to SGTA in the plant system.

Although BiFC data showed that TTOP interacted with RPN6/10/12/13, Alphafold2 only predicted that the UBL domain of TTOP could interact with the pru domain of RPN13 (Fig. 8A). This interaction was experimentally confirmed by ITC experiment (Fig. 8D). The interaction affinity (29 μM) is comparable to the affinity between human RPN13^PRU^ domain with the UBL domain of hPLC2 (9.5 μM) (Chen et al. 2016; VanderLinden et al. 2017) and the affinity between human RPN13^PRU^ domain with the UBL domain of human HR23a (26 μM) (Husnjak et al. 2008; Chen et al. 2016), and is stronger than the affinity between human RPN13^PRU^ domain with ubiquitin (91 μM) (VanderLinden et al. 2017). This mean that TTOP could have the similar 26S interaction ability like that of hPLC2 and HR23a. Bear in mind that the a.a. sequence between residues 96 – 826 of TTOP is completely novel and no structure can be predicted, it is not known if this region of TTOP could interact with the other RPN subunits. While Alphafold2 may not be able to predict all interactions precisely, it is also possible that not all BiFC signals were generated from direct interaction between TTOP and these RPN subunits (Fig. 5). It is also possible that the binding of TTOP and RPN13 may indirectly enhance the formation of complex between TTOP and the other RPN subunits. The interaction between TTOP and 26S proteosome could become stronger when more than one interacting partner is involved. Without binding to ubiquitinated substrates, 26S proteosome is inactive (Eisele et al. 2018). All seven tested UBL-containing proteins were able to stimulate peptide hydrolysis of 26S proteosome, and all three tested UBL-containing proteins were able to stimulate ATPase activities of 26S proteosome (Collins and Goldberg 2020). It will be interesting to investigate whether plant UBL-containing proteins, such as TTOP, could also activate plant 26S proteosomes.

The E1/E2/E3 ligase-CDC48A-TTOP-26S proteasome degradation pathway may also apply to the plant Tom20 receptors. In yeast, a ubiquitin-proteasome pathway for the degradation of Tom70, a TOM receptor lacking a plant ortholog (Ghifari et al. 2018), has been characterized (Wu et al. 2016). Tom70 is ubiquitinated by the cytosolic E3 ligase RSP5 (Reverses Spt-Phenotype 5) and binds to the ubiquitin binding domain of Doa1 (Degradation Of Alpha 1), which in turn binds to Cdc48 of the Cdc48/Ufd1/Npl4 complex via its C-terminal PUL domain (Mullally et al. 2006). Surprisingly, this ubiquitin-proteasome pathway is specific to Tom70, as it does not target the other yeast TOM receptor, Tom22, for degradation (Wu et al. 2016). In addition to this pathway for mitochondrial OM proteins, yeast also possesses a mitochondrial protein translocation-associated degradation (mitoTAD) pathway. The transmembrane protein Ubx2 (Ubiquitin regulatory X 2), which associates with the TOM complexes, can recruit the Cdc48/Ufd1/Npl4 complex (Schuberth and Buchberger 2005) to remove clogged precursor proteins from the TOM channel (Martensson et al. 2019). Our results revealed that there is a converged ubiquitin-proteasome degradation pathway, mediated by specific E3 ligases, CDC48 and TTOP, for tail-anchored receptors of TOC and TOM in Arabidopsis. This pathway is important for chloroplast (Ling et al. 2012) and chromoplast biogenesis (Ling et al. 2021), etiolation (Fig. 3D) and heat stress (Fig. S15). It is unclear how depletion of the components of the CHLORAD pathways, such as SP1 (Ling et al. 2012) and TTOP (Fig. 3E), affect chloroplast biogenesis. Depletion of the CHLORAD pathways may cause accumulation of the receptor components of the TOC complexes (Fig. 4K), which may affect the stoichiometry and thus the assembly of the TOC complexes. The accumulation of the TOC receptors may enhance ROS production under stresses. The CHLORAD pathway has been shown to help plants to respond to salinity and osmotic stresses by depleting TOC complexes and thus reducing ROS production from the photosystems (Ling and Jarvis 2015). Here, we showed that heat stress can induce TTOP mRNA transcription and *ttop* mutants are more sensitive to heat stress (Fig. S15). Hence, the CHLORAD pathway may also respond to heat stress and could alleviate heat stress by depleting TOC complexes and reducing heat-induced ROS. *TTOP* is also highly expressed in active sites of cell division (Fig. 4A-J), where mitochondria are abundant and active to meet the high energy demand of cell division. Nonetheless, loss of *TTOP* function did not affect plant growth under normal growth conditions. Perhaps TTOP is required only when the turnover of TOC and TOM receptors is extremely high (e.g., during etiolation) and under stresses (e.g. heat). When their turnover is low, the absence of TTOP may not cause significant phenotypes, possibly because of a lower chance of protein aggregation of TOC and TOM receptors. There is only one *TTOP* gene in the Arabidopsis genome, and no homologous protein has been identified in any non-plant species (Fig. S17). Hence, this converged ubiquitin-proteasome pathway for the degradation of tail-anchored receptors may have co-evolved with the TOC/TOM receptors during the endosymbiosis processes in plant species. While the general steps (ubiquitination by E3 ligases, retrotranslocation by CDC48, shuttling to 26S proteosomes by UBL-containing proteins, and 26S proteosome degradation) are in common, there are mechanistic differences between the animal and plant systems. In term of masking the TMD motifs of TA receptors to prevent protein aggregation during their cytosolic shuttling to 26S proteosome, plant TTOP can bind to TMDs directly, whereas animal BAG6 requires bridging to SGTA via Ubl4A for TMD binding. TTOP may evolve for direct binding to the TMDs of both TOC/TOM TA receptors to simplify the process.

## MATERIALS AND METHODS

### Plant growth conditions and generation of mutants and transgenic lines

All wild-type, mutant, and transgenic Arabidopsis (*Arabidopsis thaliana*) plants used in this study were from the Columbia-0 accession (Col-0). Plants were grown on soil or on Murashige-Skoog (MS) agar medium in petri plates at 22°C under a 16-h-light/8-h-dark cycle using cool white fluorescent light bulbs (100 µmol m^−2^ s^−1^). Seeds were surface sterilized, sown on half-strength MS medium with 1% (w/v) sucrose (pH 5.8-6.2), stratified at 4°C in the dark for 3 days, and then released in a growth chamber. To generate transgenic plants, all constructs were introduced into Agrobacterium (*Agrobacterium tumefaciens* GV3101 strain) and transformed into plants by the floral dip method (Clough and Bent 1998). The transgenic plants were selected based on resistance to the appropriate antibiotic, and then protein accumulation in the plants was confirmed by confocal microscopy or immunoblotting. Homozygous lines were obtained after propagation and selection for several generations and used for experiments. To induce the expression of *TTOP* in pTA7002-*GFP-TTOP* plants, 10 μM DEX (10 mM stock dissolved in DMSO) was added to the culture medium or 25 μM DEX was dissolved in ddH_2_O and sprayed on the plants. The equivalent volume of DMSO was added to the culture medium or ddH_2_O as negative control. The *ttop* knock-out mutants were generated using a CRISPR/Cas9 system (Pan et al. 2016). The designed single guide RNA (sgRNA) sequence CTCTAGTAGCACCAATGCGT was cloned into plasmid pKI1.0R and the plasmid transformed into Col-0 (WT) plants. The transformants were selected by antibiotic resistance. The *ttop* mutants were identified from the above selected transformants by genomic PCR and sequencing. To ensure genetic stability, Cas9-free *ttop* mutants that did not carry the CRISPR/Cas9 cassette were obtained by selecting only those seeds showing no red fluorescence under a stereomicroscope (model SZX16, Olympus). The Cas9-free seedlings were grown to maturity and their progeny used for experiments.

The *ttop-5 p35S:mCherry-Toc33* transgenic plants were generated by transforming the plasmid *p35S::mCherry-Toc33* into the *ttop-5* mutant. Homozygous *ttop-5 p35S:mCherry-Toc33* plants were obtained and crossed to WT plants, and WT transgenic plants harboring the *p35S:mCherry-Toc33* transgene were selected from the second generation of the crossed plants by sequencing of the respective PCR-amplified genomic sequences. Two individual homozygous transgenic lines expressing mCherry-Toc33 in the *ttop-5* and WT backgrounds were obtained: *ttop-5 p35S:mCherry-Toc33* #1, *ttop-5 p35S:mCherry-Toc33* #2, *p35S:mCherry-Toc33* #1, and *p35S:mCherry-Toc33* #2. Similarly, *ttop-5 p35S:mCherry-Tom20-2* transgenic plants were generated by transforming the construct *p35S:mCherry-Tom20-2* into the *ttop-5* mutant background. Homozygous *ttop-5 p35S:mCherry-Tom20-2* plants were then crossed to WT plants, and WT plants harboring the *p35S::mCherry-Tom20-2* transgene were identified. Two individual transgenic lines each in the *ttop-5* and WT background were obtained: *ttop-5 p35S:mCherry-Tom20-2* #1 and *ttop-5 p35S:mCherry-Tom20-2* #2, and *p35S:mCherry-Tom20-2* #1 and *p35S:mCherry-Tom20-2* #2.

De-etiolation experiments were performed as reported previously (Ling et al. 2012). Seeds harvested from the same growth batch were sown on MS medium, stratified at 4°C in the dark for 3 days, and then exposed to light for 6 h to induce germination. Next, the seedlings were grown in the dark for 6 days and transferred to continuous light for different periods, and their survival rates and seedling growth phenotypes were recorded. Organellar morphology in cotyledons was examined by transmission electron microscopy (TEM). Each experiment was performed independently four times, and more than 100 seedlings per genotype were used in each experiment for survival rate analysis.

To mark mitochondria in plant cells, seedlings were incubated in half-strength MS liquid culture medium containing 1 mM MitoTracker (Thermo Fisher Scientific, USA) for 30 min, and the dye was removed by washing with culture medium twice. To induce the expression of *TTOP* in pTA7002-*GFP-TTOP* transgenic plants, seedlings were grown on MS medium containing 10 μM dexamethasone (DEX) or sprayed with 25 μM DEX dissolved in ddH_2_O.

### Yeast two-hybrid, protoplast transient expression, and BiFC assays

Yeast two-hybrid analysis was performed using the MatchMaker GAL4 Two-Hybrid System 3 (Clontech, USA) according to the manufacturer’s instructions. The different CDSs were cloned into the pGBKT7 or pGADT7 vector. Pairs of pGBKT7 and pGADT7 vectors carrying the appropriate CDSs were co-transformed into the Y2HGold yeast strain. Diploids were selected on synthetic defined (SD) medium lacking tryptophan (Trp) and leucine (Leu) (SD –Trp –Leu) to select transformants. The colonies growing on SD –Trp –Leu medium were then transferred to selective SD medium lacking histidine (His), Trp, and Leu (SD –His –Trp –Leu) or lacking adenine (Ade), His, Trp, and Leu (SD –His –Ade –Trp –Leu) and were incubated at 30°C for 3–6 days to investigate protein–protein interactions. The control vector pairs pGADT7-T and pGBKT7-lam, and pGADT7-T and pGBKT7-53 were co-transformed into yeast as negative and positive controls, respectively. Yeast colonies were grown on selective SD –His –Ade –Trp –Leu medium. The experiments were repeated three times independently with similar results.

Arabidopsis mesophyll protoplast preparation and transfection were performed according to a previously described method (Yoo et al. 2007). Protoplasts were isolated from rosettes of 4-week-old Col-0 plants grown on soil by incubating the rosettes with enzyme solution containing 1% (w/v) cellulose (Onozuk R-10, Yakult, Japan) and 0.2% (w/v) macerozyme (R-10, Yakult, Japan) for 2–4 h. The isolated protoplasts were washed with W5 buffer three times to thoroughly remove the enzymes. PEG4000 was used to transfect plasmids into protoplasts; 0.1 mL (10^5^) of protoplasts was transfected with 8 μg of DNA. The transfected protoplasts were cultured for 12 h in darkness to allow protein accumulation and were collected for fluorescence detection or protein extraction. To mark the mitochondria in the cells, protoplasts were incubated in W5 culture medium containing 100 nM MitoTracker for 20 min, and excess MitoTracker was removed by washing with W5 culture medium twice. For heat treatment, the protoplasts were incubated at 37 °C for 30 min before confocal observation.

For the BiFC assay, Arabidopsis mesophyll protoplasts were co-transfected with the appropriate pairs of constructs carrying either *nY* or *cY*, fused to different CDSs, and cultured overnight (12 h) for protein expression. The reconstitution of YFP fluorescence, indicating protein–protein interaction, was detected by fluorescence microscopy using a confocal microscope. For each experiment, more than 40 individual cells were analyzed by confocal imaging that represented >90% of cells that showed similar protein expression levels and patterns. Images were captured using a Leica SP8 laser scanning confocal microscope (Leica Microsystems, German). To avoid possible cross-talk between the fluorescence channels, sequential scanning was used when necessary. Images were processed and assembled using Photoshop CS6 software (Adobe, USA).

### Statistical analysis

Statistical calculations (mean, SEM, and *t* test) were performed using Microsoft Excel software. The statistical significance of differences between two experimental groups was assessed using a two-tailed Student’s *t*-test. Differences between two datasets were considered significant at *P* < 0.05.

### Accession numbers

The Arabidopsis Genome Initiative locus identifiers for the genes mentioned in this article are as follows: *TTOP* (At5g42220), *Toc33* (At1g02280), *Toc34* (At5g05000), *Tom20-2* (At1g27390), *Tom20-3* (At3g27080), *Tom20-4* (At5g40930), *PAP2* (At1g13900), *CDC48A* (At3g09840), SP1 (At1g63900), *SP2* (At3g44160), *RPN6* (At1g29150), *RPN10* (At4g38630), *RPN12* (At1g64520), *RPN13* (At2g26590), *AtUba1* (At2g30110), *AtUbc8* (At5g41700), and *AtUb* (At4g02890).

### Data analysis

All data are presented as means with standard errors (mean ± SEM). The collected data were analyzed for statistical significance using analysis of variance (ANOVA) with Tukey’s HSD, paired *t*-tests, or unpaired *t*-tests at *P* < 0.001, *P* < 0.01, and *P* < 0.05 by SPSS (version 22).

## Data and materials availability

All data are available in the main text or the supplementary materials. Materials are available from the corresponding authors upon request.

## ACKNOWLEDGEMENTS

This work was supported by the Science, Technology and Innovation Commission of Shenzhen Municipality (Basic Research Program 201708183000803); the Hong Kong Research Grants Council Area of Excellence Scheme (AoE/M-403/16), and the Innovation and Technology Fund (Funding Support to State Key Laboratory of Agrobiotechnology) of the Hong Kong Special Administrative Region, China. Any opinions, findings, conclusions or recommendations expressed in this publication do not reflect the views of the Government of the Hong Kong Special Administrative Region or the Innovation and Technology Commission.

## AUTHOR CONTRIBUTIONS

B.L.L. and M.Y. designed the study. S.L.L., J.Y.Z., and K.C.C. generated the majority of constructs and plant transgenic lines. S.C. carried out the *in vitro* ubiquitination experiments and AlphaFold2 prediction. Z.Z. carried out phylogenetic analysis. L.Y. carried out some BiFc experiments. M.Y. generated some constructs and plant lines and carried out all the remaining experiments. B.L.L. and M.Y. wrote the manuscript. All authors revised and approved the manuscript.

