## Supplementary figures for "A converged ubiquitin-proteasome pathway for the degradation of TOC and TOM tail-anchored receptors"

**This PDF file includes:**

**Supplemental Fig. S1-17**

**Table S1 to S3**

**Supplementary Materials and Methods**

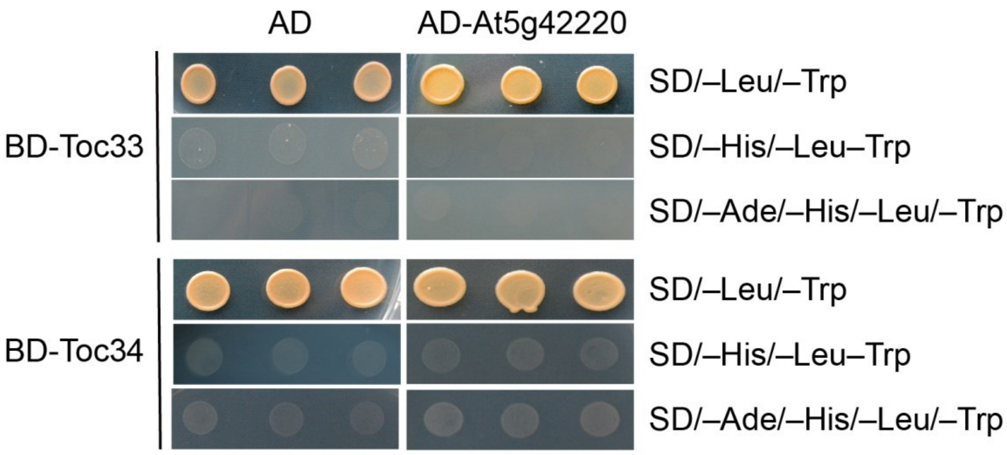


**Fig. S1. Yeast two-hybrid analysis of the interaction between TTOP and Toc33/Toc34.** Yeast was co-transformed with the indicated pairs of constructs. The growth of yeast on selective plates is shown. Growth of yeast on SD –Leu –Trp medium confirms the presence of the pair of vectors. Failure of yeast growth on SD –His –Leu –Trp and SD –Ade –His –Leu –Trp medium indicates the absence of protein–protein interaction between TTOP and Toc33/Toc34 in yeast cells.

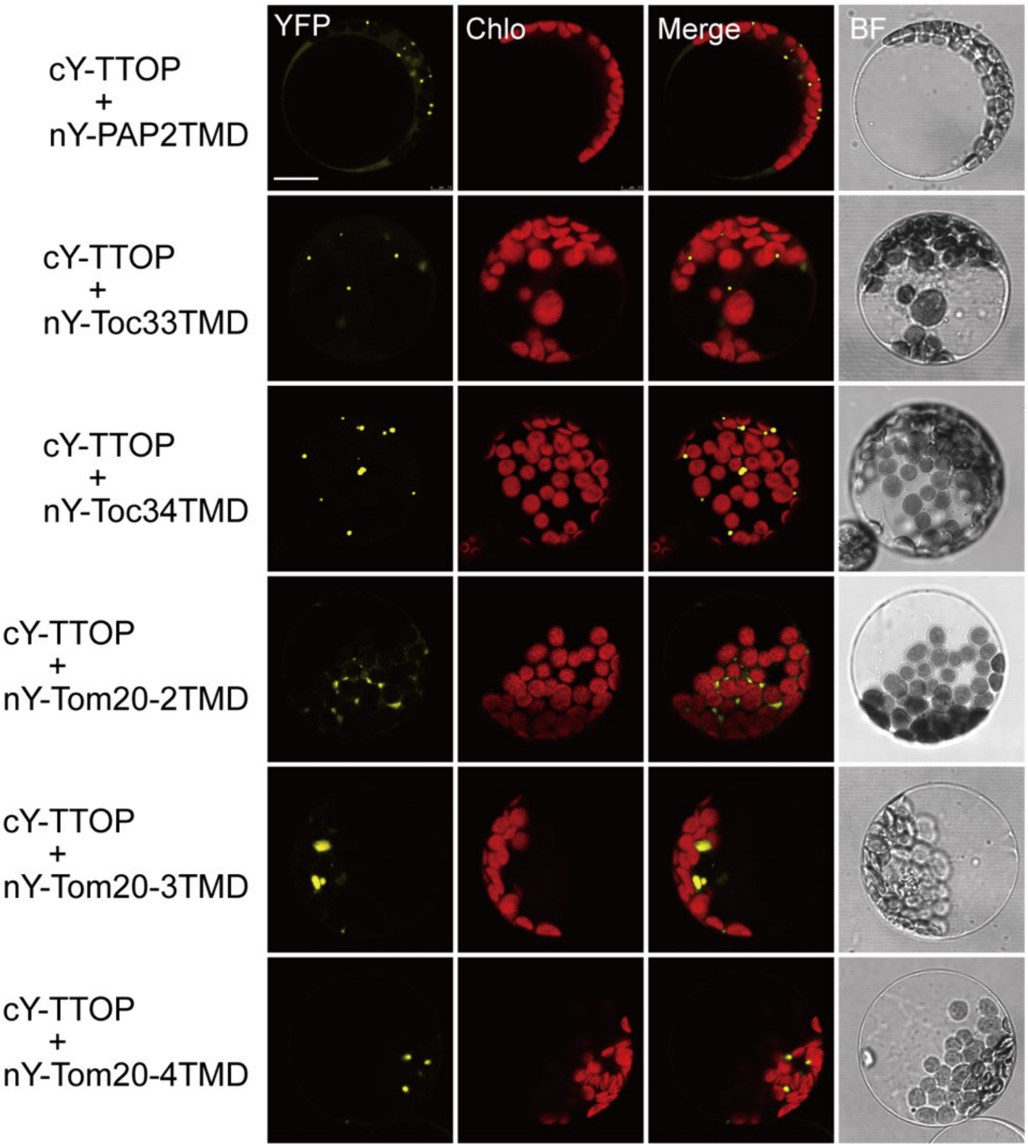


**Fig. S2. BiFC analysis of the interaction between TTOP and the TMD motifs of TA proteins.**

Protoplasts were co-transfected with the indicated pairs of constructs, which encode the fusion proteins carrying nY or cY, respectively. The TMD motifs include PAP2TMD (a.a. 615-636), Toc33TMD (a.a. 267-282), Toc34TMD (a.a. 269-287), Tom20-2TMD (a.a. 183-200), Tom20-3TMD (a.a. 175-192) and Tom20-4TMD (a.a. 162-178). Reconstitution of YFP fluorescence in protoplasts was detected by CLSM, and representative confocal images are shown. Chlo, chloroplast auto-fluorescence. BF, brightfield. Scale bar, 10 μm.

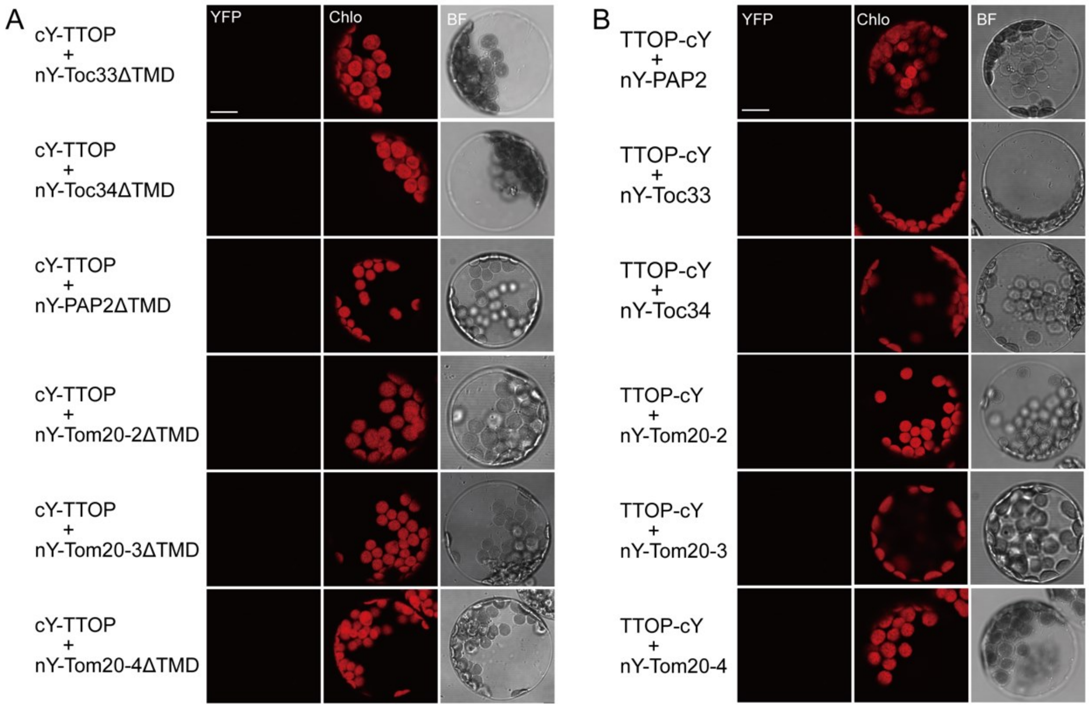


**Fig. S3. No interaction is detected between TTOP and TA proteins without TMD motifs.** Protoplasts were co-transfected with the indicated pairs of constructs encoding the fusion proteins carrying nY or cY, respectively. The TMD-truncated TA proteins include Toc33ΔTMD (a.a. 1-266 & 283-297), Toc34ΔTMD (a.a. 1-268 & 288-313), PAP2ΔTMD (a.a. 1-614 & 637-656), Tom20-2ΔTMD (a.a. 1-182 & 201-210), Tom20-3ΔTMD (a.a. 1-174 & 193-202), and Tom20-4ΔTMD (a.a. 1-161 & 179-187). Reconstitution of YFP fluorescence in protoplasts was analyzed by CLSM, and representative confocal images are shown. No interaction was detected when the TMD motifs of the TA proteins were truncated **(A)** or when cY was fused to the C terminus of TTOP **(B)**. Chlo, chloroplast auto-fluorescence. BF, brightfield. Scale bar, 10 µm.

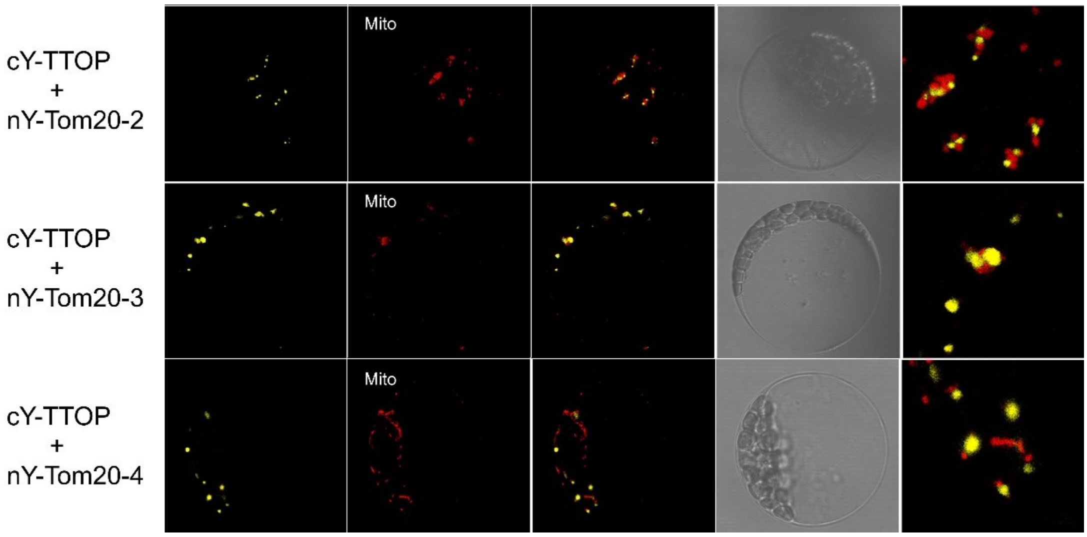


**Fig. S4. Bimolecular fluorescence complementation (BiFC) analysis of the interaction between TTOP and Tom20 proteins.** Protoplasts were transiently co-transfected with the indicated pairs of constructs, which encode the fusion proteins carrying complementary N- or C-terminal YFP fragments (nY or cY), respectively. Reconstitution of YFP fluorescence in protoplasts was detected by confocal microscopy; representative images are shown. Mito, mitochondria marked with MitoTracker. BF, brightfield. Scale bar, 10 µm.

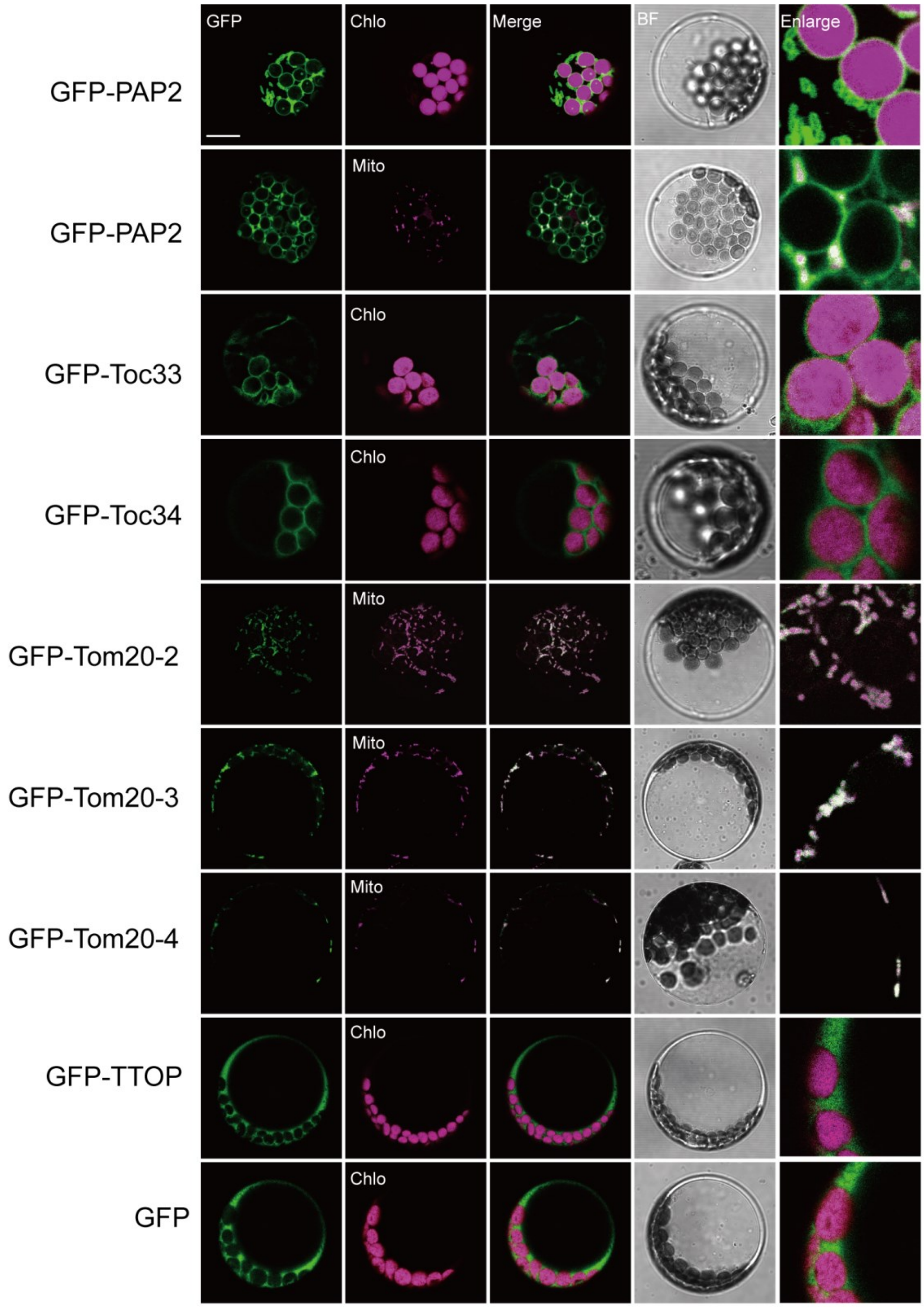


**Fig. S5. Subcellular localization of TTOP and chloroplast and mitochondrial outer membrane TA proteins in protoplasts.**

Protoplasts were transfected with the indicated constructs encoding fusion proteins carrying a GFP protein at their N terminus. The GFP fluorescence in protoplasts was analyzed by CLSM, and representative confocal images are shown. A construct encoding free GFP was also transfected in protoplasts as a positive control for GFP fluorescence and as a negative control for organellar localization. Chlo, chloroplast auto-fluorescence; Mito, mitochondria marked with MitoTracker. BF, brightfield. Scale bar, 10 µm.

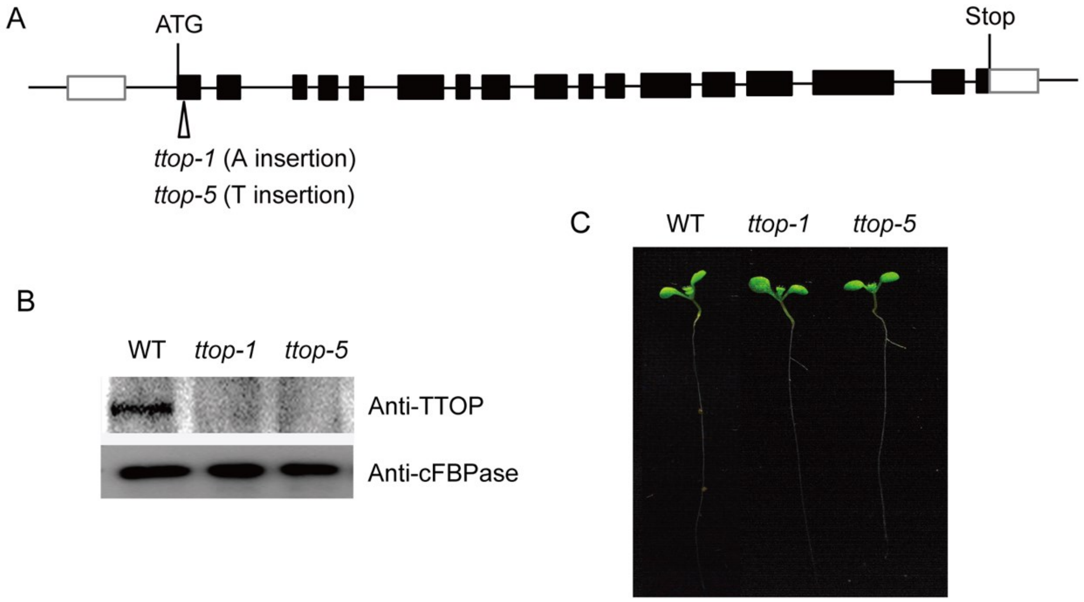


**Fig. S6. Identification and growth phenotype of *ttop* mutant seedlings.**
**(A)** Schematic representation of the *TTOP* (At5g42220) locus, annotated with the positions of the *ttop-1* and *ttop-5* mutations. ttop-1 and ttop-5 has a single-base insertion of an A or T in the 43 bp of the first TTOP exon, respectively. Black boxes, exons; white boxes, untranslated regions; black lines, introns. **(B)** Immunoblotting analysis of TTOP abundance in the wild-type Col-0 (WT) and *ttop1-1* and *ttop1-5* seedlings with the rabbit anti-TTOP antibody developed in this study. Anti-cFBPase was used as a loading control. **(C)** Representative growth phenotypes of 8-day-old WT, *ttop1-1*, and *ttop1-5* seedlings grow under normal conditions.

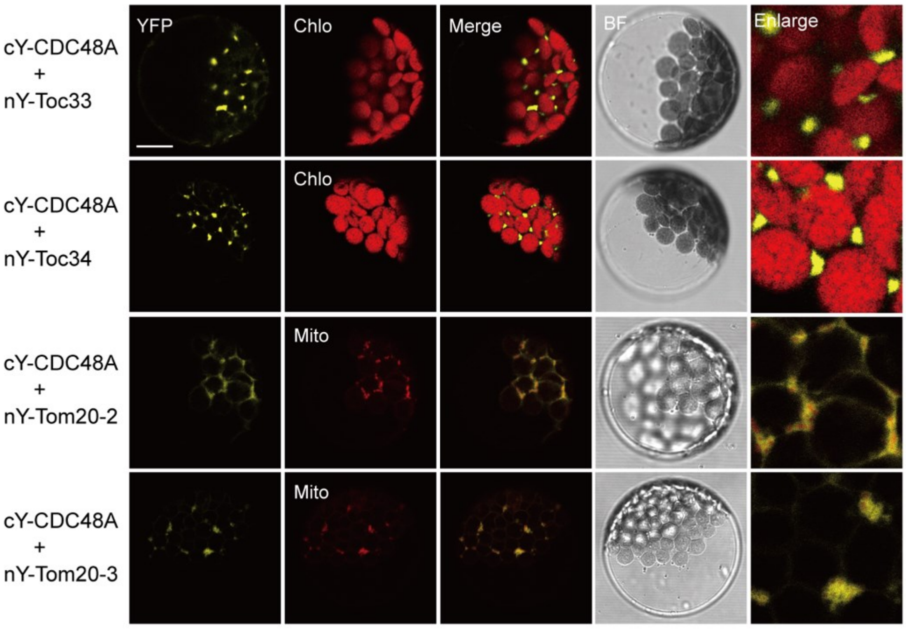


**Fig. S7. BiFC analysis of the interaction between CDC48A and TA proteins in the absence of TTOP.**

Protoplasts were prepared from the *ttop1-5* mutant and were co-transfected with the indicated pairs of constructs encoding CDC48A or TA proteins carrying nY or cY in their respective N termini. Reconstitution of YFP fluorescence in protoplasts was detected by CLSM; representative confocal images are shown. Chlo, chloroplast auto-fluorescence; Mito, mitochondria marked with MitoTracker. BF, brightfield. Scale bar, 10 µm.

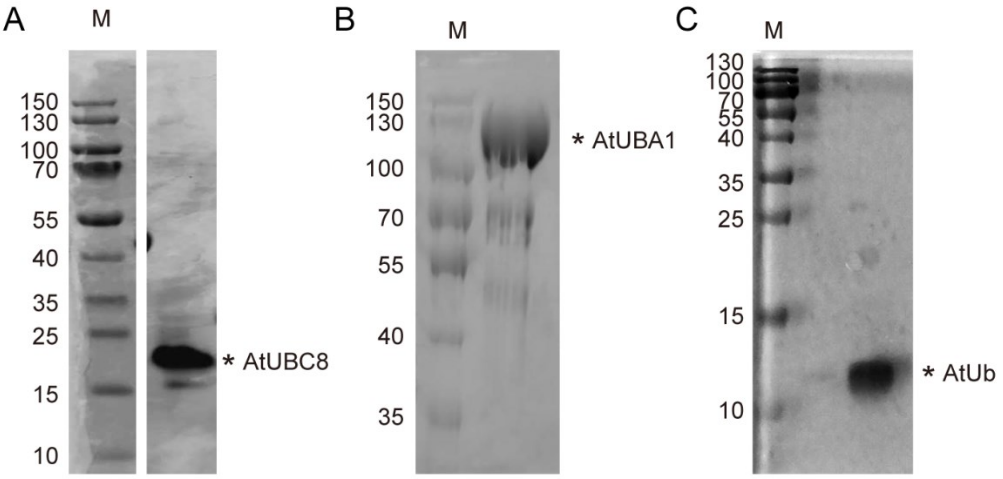


**Fig. S8. Purification of recombinant AtUBC8, AtUBA1, and AtUb.**

SDS-PAGE analysis of purified AtUBC8 **(A)**, AtUBA1 **(B)** and AtUb (**C)** proteins after removal of GST-tag and purification by gel-filtration. Positions of molecular weight markers are indicated at left (sizes in kDa).

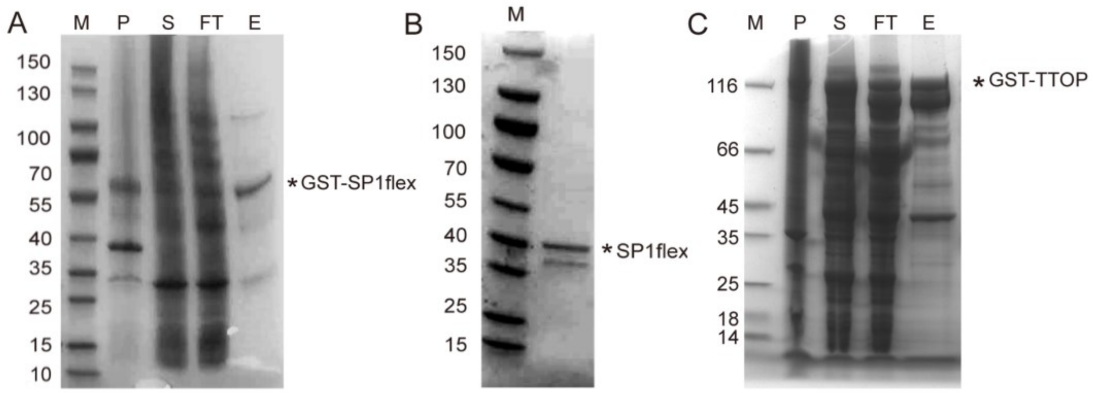


**Fig. S9. Purification of recombinant SP1flex and TTOP.**
**(A)** SDS-PAGE analysis of GST-SP1flex protein purified using a GST column. **(B)** SP1flex purified by gel filtration after PreScission Protease digestion of GST-SP1flex. **(C)** SDS-PAGE analysis of GST-TTOP protein purified using a GST column. Positions of molecular weight markers are indicated to the left (sizes in kDa). M, protein marker; P, pellet; S, supernatant; FT, flow-through; E, eluent. BD, before protease digestion. AD, after protease digestion. GF, gel filtration.

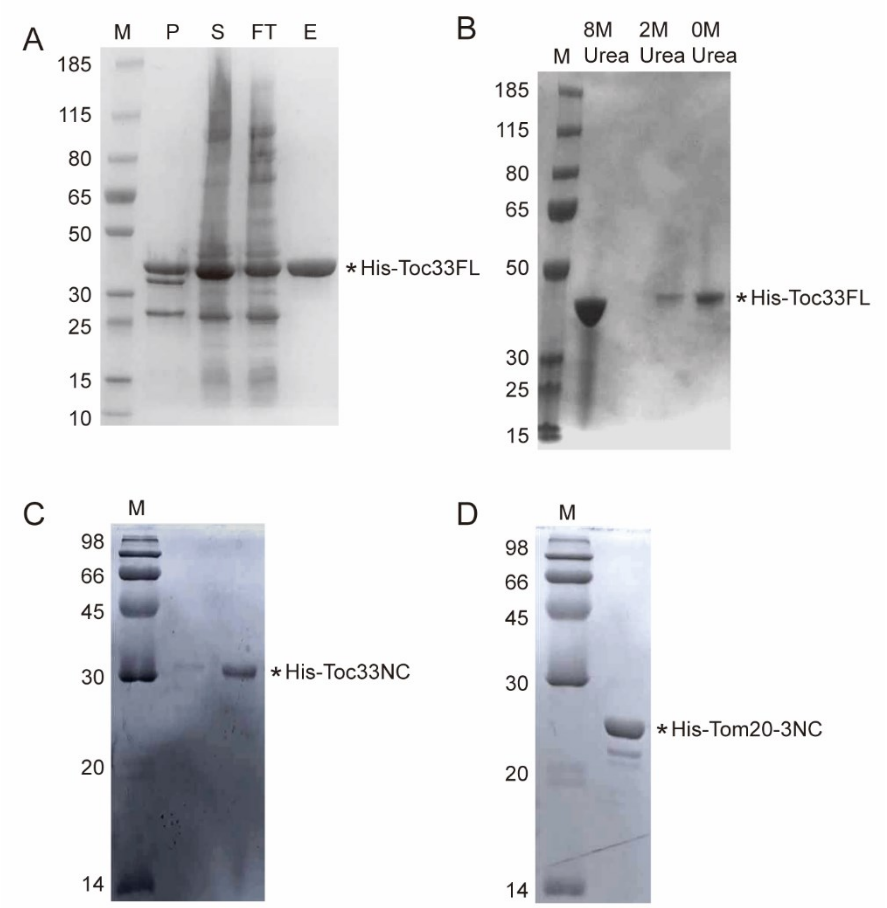


**Fig. S10. Purification of recombinant Toc33FL, Toc33NC, and Tom20-3NC.**
**(A)** SDS-PAGE analysis of unfolded His-Toc33FL purified using a Ni column. **(B)** Refolding of His-Toc33FL protein upon removal of urea by dialysis. **(C** and **D)** SDS-PAGE analysis of purified His-Toc33NC **(C)** and His-Tom20-3NC **(D)** proteins using a Ni column. Positions of molecular weight markers are indicated at left (sizes in kDa). M, protein marker; P, pellet; S, supernatant; FT, flow-through; E, eluent.

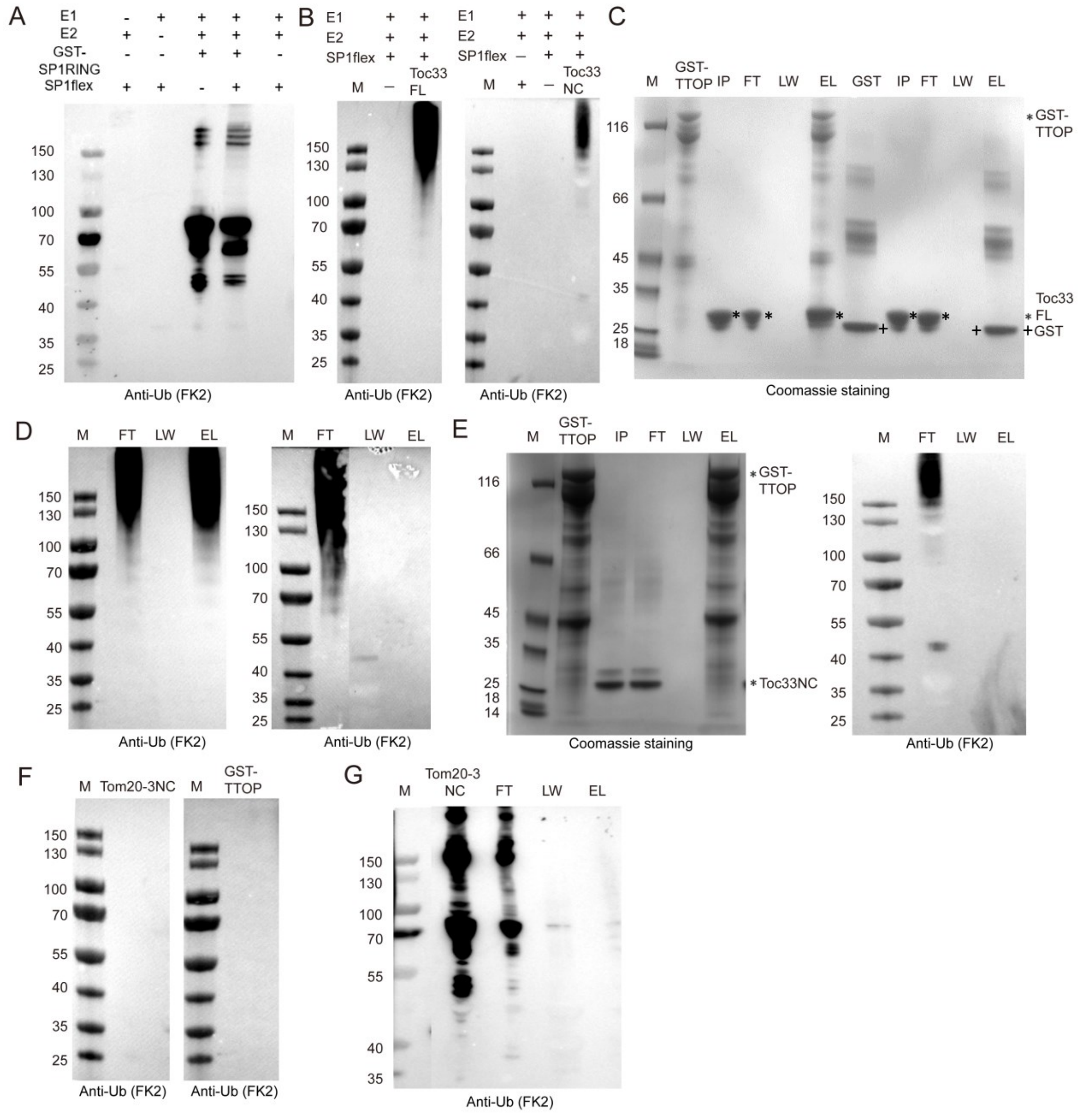


**Fig. S11.** ***In vitro* ubiquitination assays of tail-anchored receptors.**

**(A)** Analysis of SP1 auto-ubiquitination activity *in vitro*. Recombinant SP1flex was purified and tested for self-ubiquitination activity. Western blotting was analyzed with anti-ubiquitin (FK2, MERCK). SP1flex cannot self-ubiquitinate (lane 6), but only GST-SP1RING (lane 4) and GST-SP1 RING plus SP1flex (lane 5) can self-ubiquitinate at the GST fusion protein. **(B)** *In vitro* ubiquitination of His-Toc33FL (a.a. 1-297) and His-Toc33NC (a.a. 1-266) by recombinant SP1flex. **(C)** GST-TTOP but not GST alone can pull down non-ubiquitinated His-Toc33FL. IP: input His-Toc33FL; FT: Flow through; LW; last washing fraction; EL: eluted proteins. (**D)** GST-TTOP (left panel) but not GST alone (right panel) can pull down ubiquitinated His-Toc33FL. (**E**) Neither non-ubiquitinated (left panel) nor ubiquitinated (right panel) His-Toc33NC was pulled down by GST-TTOP. **(F)** His-Tom20-3NC (left panel) and TTOP (right panel) were not ubiquitinated by recombinant SP1flex and UBC8 *in vitro*. (**G**) Ubiquitination of His-Tom20-3NC (a.a. 1-174) was facilitated by plant extracts (lane 1), but ubiquitinated His-Tom20NC cannot be pulled down by recombinant GST-TTOP (lane 4). Positions of molecular weight markers are indicated at left (sizes in kDa). M, protein marker.

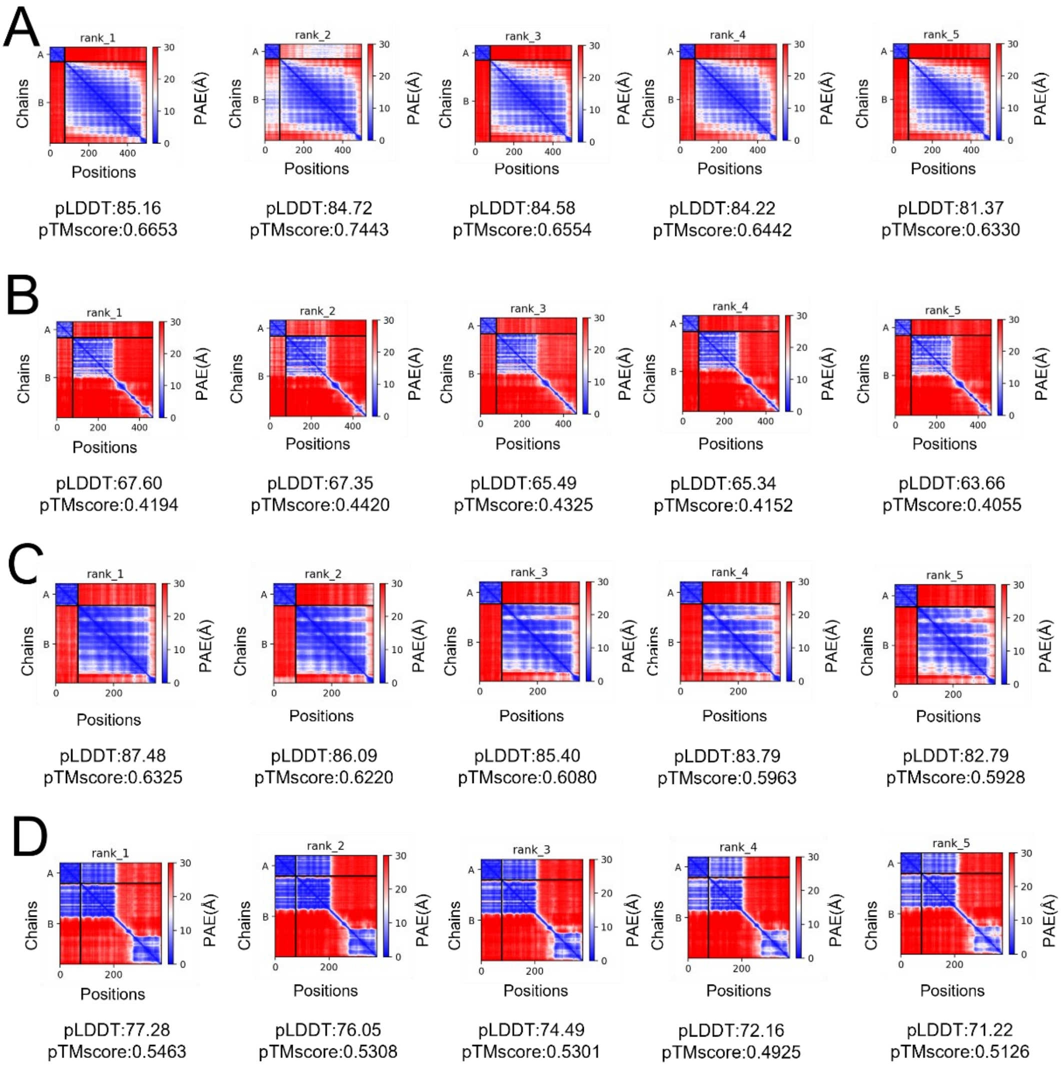


**Fig. S12. Prediction of TTOP interacting domain to RPN subunits by Alphafold**

By Alphafold2 prediction, the UBL domain of TTOP could not interact with RPN6 (A), RPN10 (B), nor RPN12 (C). It is predicted that the UBL domain of TTOP could interact with RPN13 (D).

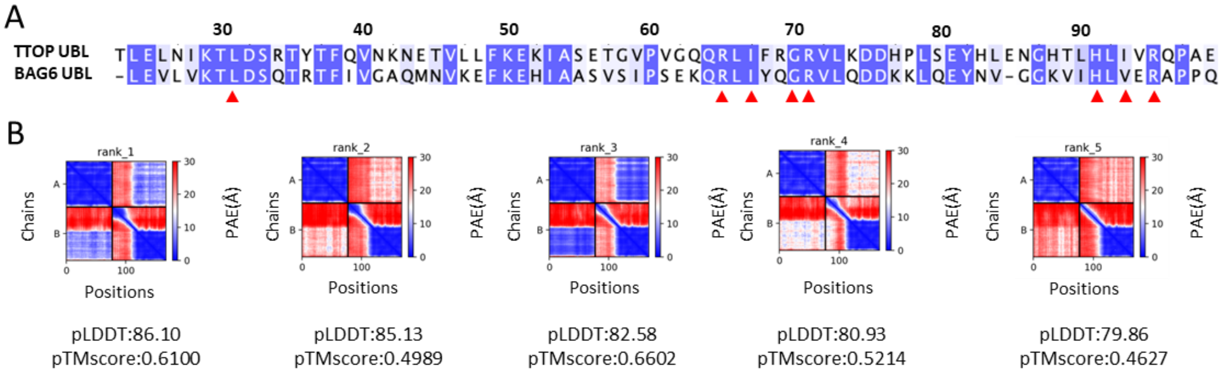


**Fig. S13. Prediction of TTOP UBL domain and SP1 RING interaction.**

Amino acid sequence alignment between the TTOP UBL and human BAG6 UBL domains. TTOP UBL contains the conserved RNF126_NZF interacting residues (Red triangle) (A). By AlphaFold2 prediction, the UBL domain of TTOP could interact with SP1 RING domain, especially rank_1 and rank_3 give much lower PAE values (B).

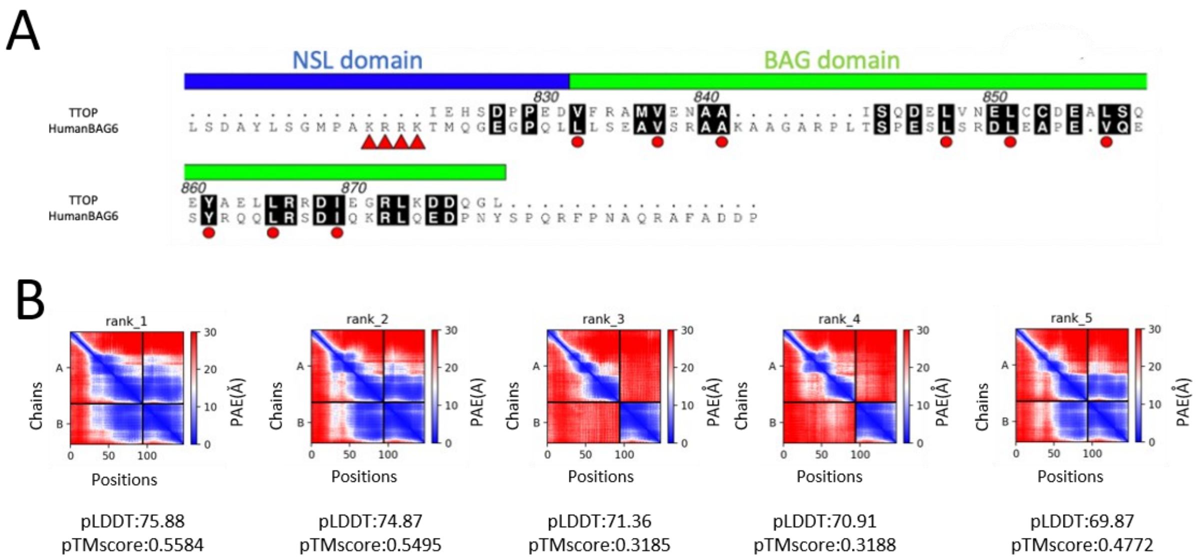


**Fig. S14. TTOP C-terminus contains a BAG domain**

Amino acid sequence alignment between the C-termini of TTOP and human BAG6. TTOP C-terminus does not contain the TRC35 interaction residues (Red triangle) in the NSL domain of BAG6 but contains a BAG domain. Especially, the TTOP C-terminal contains the conserved Ubl4A interaction residues (Red circle) (A). By AlphaFold2 prediction, the low PAE value predicted that the C-terminal of TTOP could interact with human Ubl4A (B).

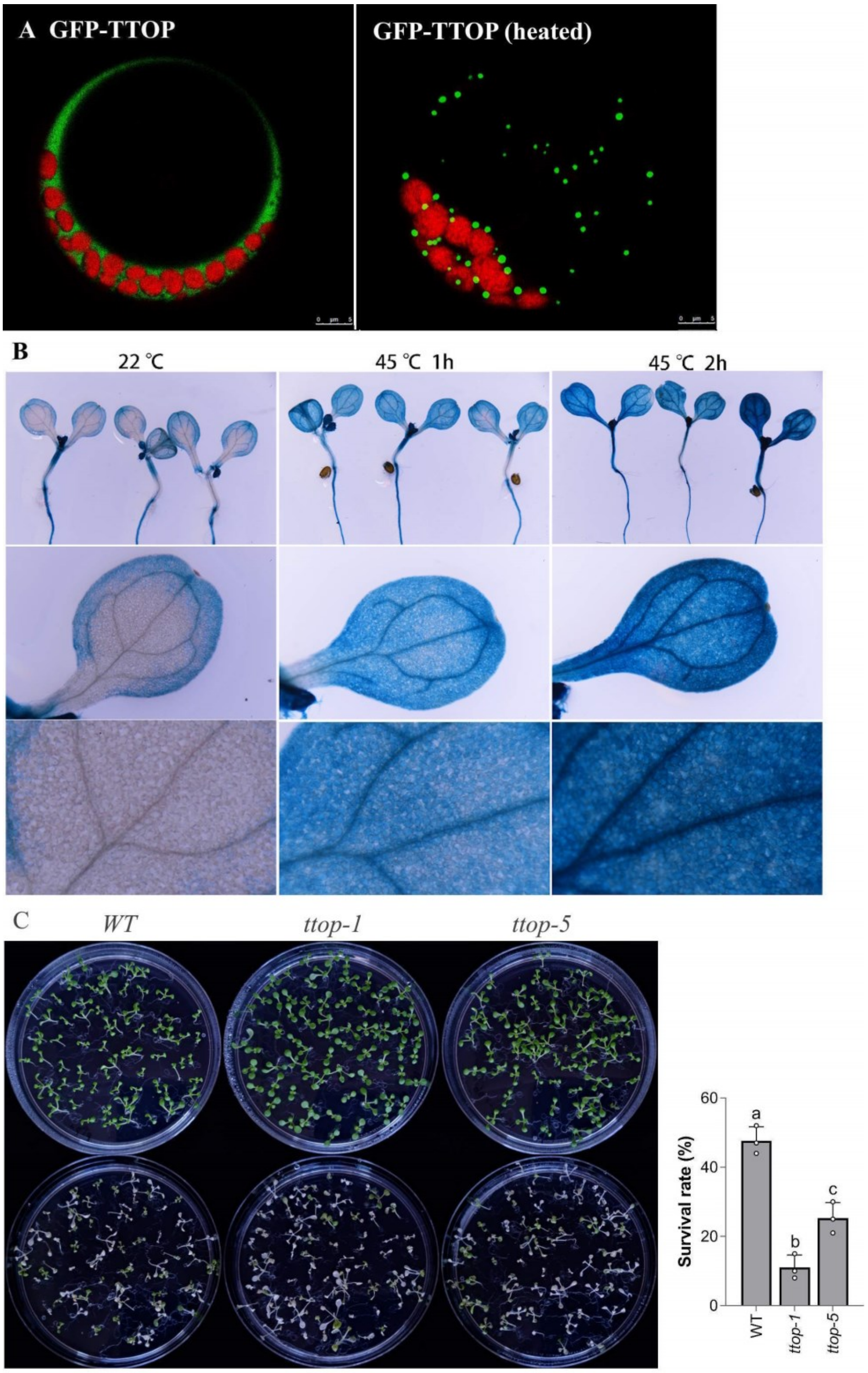


**Fig. S15. Heat treatment induces TTOP mRNA transcription.**

**(A)** Protoplasts transformed with GFP-TTOP before (leaf panel) and after subject to heat treatment at 37C for 30 mins (right panel).

**(B)** Ten-day-old *pTTOP:GUS* seedlings were used for GUS staining. For heat treatment, the seedlings were placed in 45°C incubator for 1 or 2h before GUS staining. Scale bar in A, E, I, 2.0 mm. Scale bar in B, C, D, F, G, H, 200 µm.

**(C)** Nine-day-old seedlings (upper panel) were placed in 45 °C water bath for 1 hr and then were allowed to recover at RT for 5 days (lower panel).

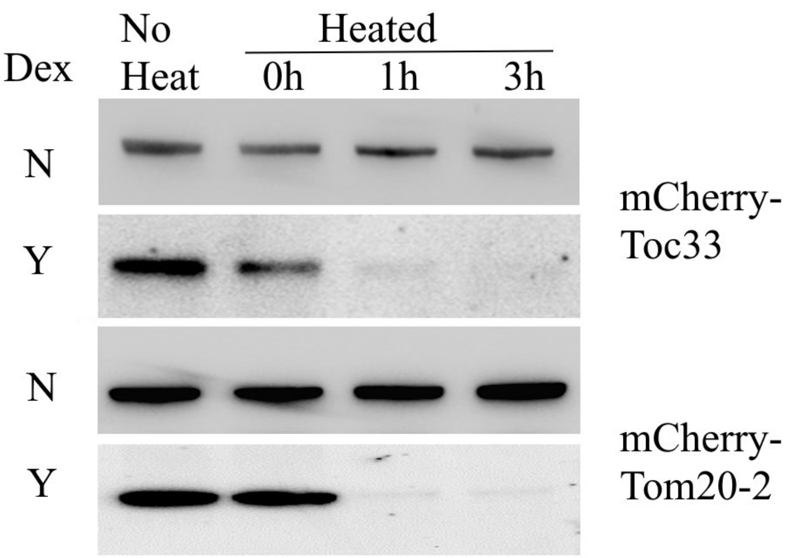


**Fig. S16. TTOP assists the degradation of mCherry-Toc33 and mCherry-Tom20-2 in planta.**

The transgenic DEX-GFP-TTOP line was transformed with 35S: mCherry-Toc33 and 35S: mCherry-Tom20-2. 25μM DEX were sprayed to the seedlings at 8-day-old to induce TTOP expression. Some seedlings were treated with DMSO as negative controls. After 24 h, seedlings were placed in 45°C incubator for 30 minutes (heated) or kept in R.T. (no heat). Heat-treated seedlings were allowed to recover in R.T. for 1 and 3 hours prior to immunoblotting with anti-mCherry antibodies.

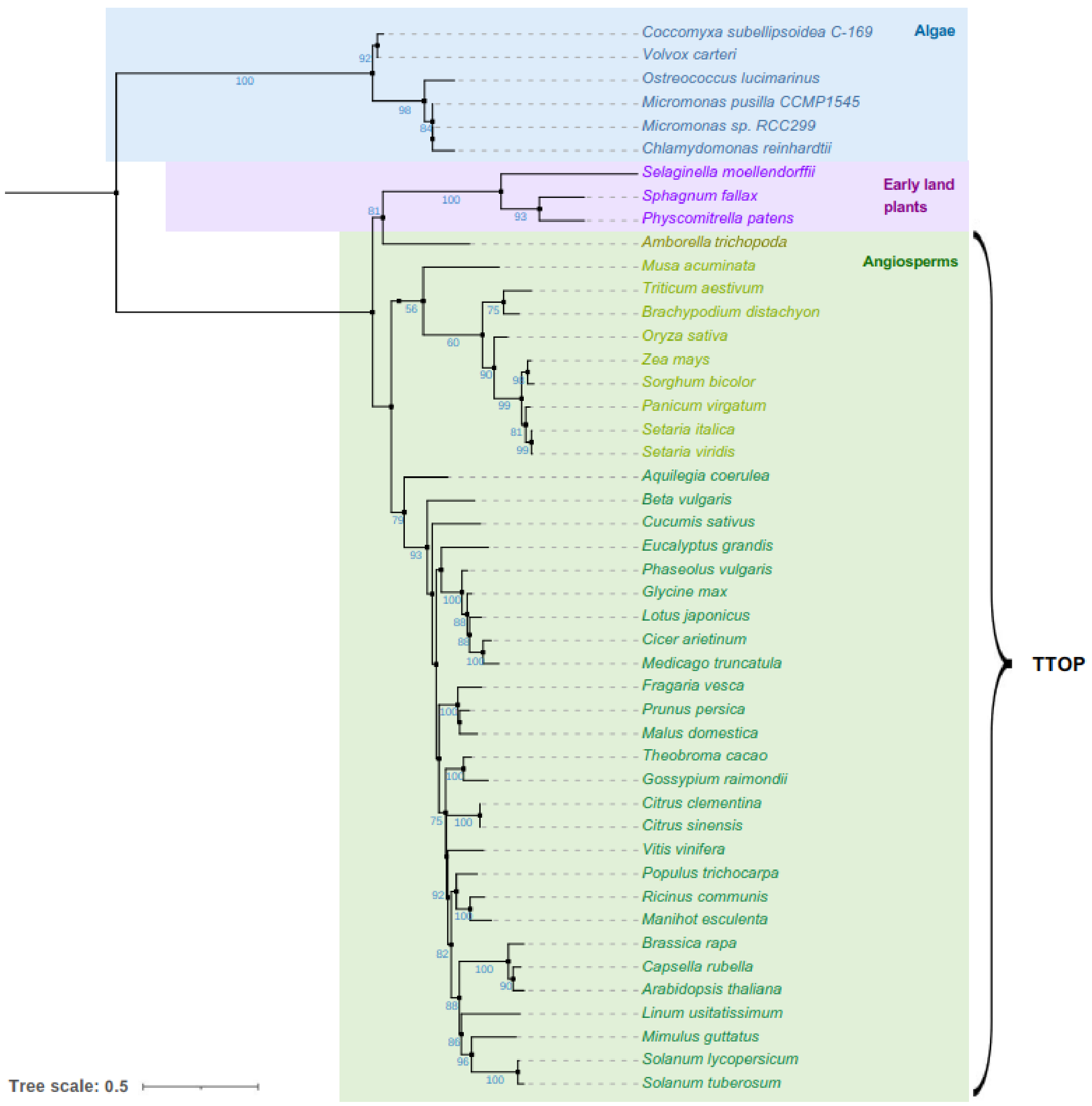


**Fig. S17. Phylogenetic tree of TTOP**

Phylogenetic analysis of TTOP was conducted using the maximum likelihood method. Percent support values from 1,000 bootstrap samples and scale bar are shown.

**Table S1. PCR primers for vector construction.**

| Primer name | Primer sequence (5′ to 3′) | Application |
|---|---|---|
| TTOP-CDS-1F | CCAGGCCTACTAGTGGATCCATGGAAGATCAACCCATTAA | Cloning *TTOP* CDS into pBI221 vector carrying *nY* or *eGFP* or *mCherry* |
| TTOP-CDS-1R | ACCCGGGAGCGGTACCTTATAGACCTTGATCATCTTTTAG | |
| TTOP-CDS-2F | TGGGCATCGATACGGGATGGAAGATCAACCCATTAA | Cloning *TTOP* CDS into pGADT7 vector |
| TTOP-CDS-2R | CAGCTCGAGCTCGATGTTATAGACCTTGATCATCTTTTAG | |
| Myc-TTOP-CDS-F | AGAACACGGGGGACTCTAGAATGGAGCAAAAGTTGATTTCTGAGGAGGATCTTATGGAAGATCAACCCATTAA | Cloning *TTOP* CDS with Myc tag in the N terminus into pBI221 vector |
| GFP-TTOP CDS-1F | TAGTCGACTCTAGCCTCGAGATGGTGAGCAAGGGCGA | Cloning *TTOP* CDS as GFP fusion in the N terminus into pTA7002 vector |
| GFP-TTOP CDS-1R | GTCGATCCTCTAGCCTCGAGTTATAGACCTTGATCATCTTTT | |
| mCherry-TTOP CDS-F | TAGTCGACTCTAGCCTCGAGATGTTGAGCAAGGGCGAG | Cloning *TTOP* CDS with *mCherry* fusion in the N terminus into pTA7002 vector |
| mCherry-TTOP CDS-R | GTCGATCCTCTAGCCTCGAGTTATAGACCTTGATCATCTTTT | |
| GFP-TTOP CDS-2F | ACTCTAGAGGATCCCCGGGATGGTGAGCAAGGGCGA | Cloning *TTOP* CDS with GFP *fusion* in the N terminus into pBI121 vector to replace *GUS* |
| GFP-TTOP CDS-2R | ATCGGGGAAATTCGAGCTCTTATAGACCTTGATCATCTTTT | |
| TTOP Pro-F | GACCATGATTACGCCAAGCTTCTTATTTACAATGGCTGCCACTGG | Cloning *TTOP* promoter into pBI121 vector carrying *GUS* to replace *35S* promoter |
| TTOP Pro-R | AGGGACTGACCACCCGGGGAACCCTAAAGTATTAAAATT | |
| PAP2-CDS-1F | CCAGGCCTACTAGTGGATCCATGATCGTTAATTTCTCTTT | Cloning *PAP2* CDS into pBI221 vector carrying *nY* or *eGFP* in the N terminus |
| PAP2-CDS-1R | ACCCGGGAGCGGTACCTTATGTCTCCTCGTTCTTG | |
| PAP2-CDS-2F | TATGGCCATGGAGGCCGAATTCATGATCGTTAATTTCTCTTT | Cloning *PAP2* CDS into pGBKT7 vector |
| PAP2-CDS-2R | TTATGCGGCCGCTGCAGGTTATGTCTCCTCGTTCTTG | |
| PAP2-CDS-3F | TTGCTCCGTGGATCCGGTACCATGATCGTTAATTTCTCTTT | Cloning *PAP2* CDS into HBT vector carrying *FLAG* tag in the C terminus |
| PAP2-CDS-3R | CTTGTAGTCAGAAGGCCTCCCGGGTGTCTCCTCGTTCTTGACTGG | |

**Table S1 (continued). PCR primers for vector construction.**

| | | |
|---|---|---|
| PAP2-ΔTMD-1R | ACCCGGGAGCGGTACCTTATGTCTCCTCGTTCTTGACTGGGATCCAACGGTTTCCAGACGAAGATTTCTTTCCCCGGGTAGCATTAGATTCTGATTTTCCC | Cloning TMD-truncated *PAP2* CDS into pBI221 vector carrying *nY* in the N terminus |
| PAP2-ΔTMD-2R | TTATGCGGCCGCTGCAGGTTATGTCTCCTCGTTCTTGACTGGGATCCAACGGTTTCCAGACGAAGATTTCTTTCCCCGGGT AGCATTAGATTCTGATTTTCCC | Cloning TMD-truncated *PAP2* CDS into in pGBKT7 vector |
| PAP2-TMD-F | GATCCTTGTGGTATGCCAAAGGAGCAGGCTTGATGGTTGTGGGTGTGCTTTTAGGGTTCATTATCGGGTTTTTTTAAGGTAC | Cloning TMD of *PAP2* into in pBI221 vector carrying *nY* in the N terminus |
| PAP2-TMD-R | CTTAAAAAAACCCGATAATGAACCCTAAAAGCACACCCACAACCATCAAGCCTGCTCCTTTGGCATACCACAAG | |
| Toc33-CDS-1F | CCAGGCCTACTAGTGGATCCATGGGGTCTCTCGTTCG | Cloning *Toc33* CDS into pBI221 vector carrying *nY* or *eGFP* in the N terminus |
| Toc33-CDS-1R | ACCCGGGAGCGGTACCTTAAAGTGGCTTTCCACTTG | |
| Toc33-CDS-2F | TATGGCCATGGAGGCCGAATTCATGGGGTCTCTCGTTCG | Cloning *Toc33* CDS into pGBKT7 vector |
| Toc33-CDS-2R | TTATGCGGCCGCTGCAGTTAAAGTGGCTTTCCACTTG | |
| Toc33-CDS-3F | TTGCTCCGTGGATCCGGTACCATGGGGTCTCTCGTTCG | Cloning *Toc33* CDS into HBT vector carrying *FLAG* tag in the C terminus |
| Toc33-CDS-3R | CTTGTAGTCAGAAGGCCTCCCGGGAAGTGGCTTTCCACTTGTC | |
| Toc33-ΔTMD-R | ACCCGGGAGCGGTACCTTAAAGTGGCTTTCCACTTGTCTTGATATCATTTCTGATTGCTCCTTG TTTCTTTCCTTTATCATCAGAG | Cloning TMD-truncated *Toc33* CDS into pBI221 vector carrying *nY* in the N terminus |
| Toc33-TMD-F | GATCCCTCATCCCTCTTATCATAGGCGCTCAGTATTTGATCGTTAAGATGATTTAAGGTAC | Cloning TMD of *Toc33* into in pBI221 vector carrying *nY* in the N terminus |
| Toc33-TMD-R | CTTAAATCATCTTAACGATCAAATACTGAGCGCCTATGATAAGAGGGATGAGG | |
| mCherry-Toc33-CDS-F | GAGGATCCCCGGGTACCATGGTGAGCAAGGGCGA | Cloning *Toc33* CDS fused to *mCherry* in the N terminus into pBI121 vector |
| mCherry-Toc33-CDS-R | GGGGAAATTCGAGCTCTTAAAGTGGCTTTCCACTTG | |
| Toc34-CDS-1F | CCAGGCCTACTAGTGGATCCATGGCAGCTTTGCAAAC | Cloning *Toc34* CDS into pBI221 vector carrying *nY* or *eGFP* in the N terminus |
| Toc34-CDS-1R | ACCCGGGAGCGGTACCTCAAGACCTTCGACTTG | |

**Table S1 (continued). PCR primers for vector construction.**

| Primer name | Primer sequence (5′ to 3′) | Application |
|---|---|---|
| Toc34-CDS-2F | TATGGCCATGGAGGCCGAATTCATGGCAGCTTTGCAAAC | Cloning *Toc34* CDS into pGBKT7 vector |
| Toc34-CDS-2R | TTATGCGGCCGCTGCAGTCAAGACCTTCGACTTGC | |
| Toc34-CDS-3F | TTGCTCCGTGGATCCGGTACCATGGCAGCTTTGCAAAC | Cloning *Toc34* CDS into HBT vector carrying *FLAG* tag in the C terminus |
| Toc34-CDS-3R | CTTGTAGTCAGAAGGCCTCCCGGGAGACCTTCGACTTGCTA | |
| Toc34-ΔTMD -R | ACCCGGGAGCGGTACCTCAAGACCTTCGACTTGCTAAACCGGAGTCTCGCAGCTCCCACGCCGGTTTACTCTCTCTCGAAACATCGGACTTGATTGCTTTTTTTCCTCTTTCGTTTGGG | Cloning TMD-truncated *Toc34* CDS into pBI221 vector carrying *nY* in the N terminus |
| Toc34-TMD -F | GATCCCTGATTCCTTTAATGTTTGCATTCCAATACTTGCTGGTGATGAAGCCATTGGTTCGATAAGGTAC | Cloning TMD of *Toc34* into in pBI221 vector carrying *nY* in the N terminus |
| Toc34-TMD -R | CTTATCGAACCAATGGCTTCATCACCAGCAAGTATTGGAATGCAAACATTAAAGGAATCAGG | |
| Tom20-2-CDS-1F | CCAGGCCTACTAGTGGATCCATGGAGTTCTCTACCGC | Cloning *Tom20-2* CDS into pBI221 vector carrying *nY* or *eGFP* in the N terminus |
| Tom20-2-CDS-1R | ACCCGGGAGCGGTACCTTATCTGGCAGGAGGTG | |
| Tom20-2-CDS-2F | TATGGCCATGGAGGCCGAATTCATGGAGTTCTCTACCGC | Cloning *Tom20-2* CDS into pGBKT7 vector |
| Tom20-2-CDS-2R | TTATGCGGCCGCTGCAGTTATCTGGCAGGAGGTG | |
| Tom20-2-CDS-3F | TTGCTCCGTGGATCCGGTACCATGGAGTTCTCTACCGC | Cloning *Tom20-2* CDS into HBT vector carrying *FLAG* tag in the C terminus |
| Tom20-2-CDS-3R | CTTGTAGTCAGAAGGCCTCCCGGGTCTGGCAGGAGGTGGAGGGCCAAG | |
| Tom20-2-ΔTMD -1R | ACCCGGGAGCGGTACCTTATCTGGCAGGAGGTGGAGGGCCAAGGGATTTATCATAAGTGAATTCAGTGTTCCTC | Cloning TMD-truncated *Tom20-2* CDS into pBI221 vector carrying *nY* in the N terminus |
| Tom20-2-ΔTMD -2R | TTATGCGGCCGCTGCAGGTTATCTGGCAGGAGGTGGAGGGCCAAGGGATTTATCATAAGTGAATTCAGTGTTCCTC | Cloning TMD-truncated *Tom20-2* CDS into pGBKT7 vector |
| Tom20-2-TMD-F | GATCCGTATGCGGTTGGATAATTCTCGCTTGTGGGATTGTTGCTTGGGTTGGCATGGCATAAGGTAC | Cloning TMD of *Tom20-2* into in pBI221 vector carrying *nY* in the N terminus |
| Tom20-2-TMD-R | CTTATGCCATGCCAACCCAAGCAACAATCCCACAAGCGAGAATTATCCAACCGCATACG | |

**Table S1 (continued). PCR primers for vector construction.**

| Primer name | Primer sequence (5′ to 3′) | Application |
|---|---|---|
| mCherry-Tom20-2-CDS-F | GAGGATCCCCGGGTACCATGGTGAGCAAGGGCGA | Cloning *Tom20-2* CDS fused to *mCherry* in the N terminus into pBI121 vector |
| mCherry-Tom20-2-CDS-R | GGGGAAATTCGAGCTCTTATCTGGCAGGAGGTG | |
| Tom20-3-CDS-1F | CCAGGCCTACTAGTGGATCCATGGATACGGAAACTGAGTT | Cloning *Tom20-3* CDS into pBI221 vector carrying *nY* or *eGFP* in the N terminus |
| Tom20-3-CDS-1R | ACCCGGGAGCGGTACCTTAACGAGGAGGAGAGACA | |
| Tom20-3-CDS-2F | TATGGCCATGGAGGCCGAATTCATGGATACGGAAACTGAGTT | Cloning *Tom20-3* CDS into pGBKT7 vector |
| Tom20-3-CDS-2R | TTATGCGGCCGCTGCAGTTAACGAGGAGGAGAGACA | |
| Tom20-3-CDS-3F | TTGCTCCGTGGATCCGGTACCATGGATACGGAAACTGAGTT | Cloning *Tom20-3* CDS into HBT vector carrying *FLAG* tag in the C terminus |
| Tom20-3-CDS-3R | CTTGTAGTCAGAAGGCCTCCCGGGACGAGGAGGAGAGACA | |
| Tom20-3-ΔTMD -1R | ACCCGGGAGCGGTACCTTAACGAGGAGGAGAGACAGGCACATTAGCTTTATCATACTTGGCATCACTACTT | Cloning TMD-truncated *Tom20-3* CDS into pBI221 vector carrying *nY* in the N terminus |
| Tom20-3-ΔTMD -2R | TTATGCGGCCGCTGCAGGTTAACGAGGAGGAGAGACAGGCACATTAGCTTT ATCATACTTGGCATCACTACTT | Cloning TMD-truncated *Tom20-3* CDS into pGBKT7 vector |
| Tom20-3-TMD -F | GATCCGCTATGGGTTGGGTGATTCTAGCCATTGGTGTTGTTGCTTGGATCAGTTTCGCGTAAGGTAC | Cloning TMD of *Tom20-3* into in pBI221 vector carrying *nY* in the N terminus |
| Tom20-3-TMD -R | CTTACGCGAAACTGATCCAAGCAACAACACCAATGGCTAGAATCACCCAACCCATAGCG | |
| Tom20-4-CDS-1F | CCAGGCCTACTAGTGGATCCATGGATATGCAGAATGAAA | Cloning *Tom20-4* CDS into pBI221 vector carrying *nY* or *eGFP* in the N terminus |
| Tom20-4-CDS-1R | ACCCGGGAGCGGTACCTTACTGCCTTGACACCG | |
| Tom20-4-CDS-2F | TATGGCCATGGAGGCCGAATTCATGGATATGCAGAATGAAAACG | Cloning *Tom20-4* CDS into pGBKT7 vector |
| Tom20-4-CDS-2R | TTATGCGGCCGCTGCAGTTACTGCCTTGACACCG | |
| Tom20-4-CDS-3F | TTGCTCCGTGGATCCGGTACCATGGATATGCAGAATGAAA | Cloning *Tom20-4* CDS into HBT vector carrying *FLAG* tag in the C terminus |
| Tom20-4-CDS-3R | CTTGTAGTCAGAAGGCCTCCCGGGCTGCCTTGACACCGGCGT | |
| Tom20-4-ΔTMD -1R | ACCCGGGAGCGGTACCTTACTGCCTTGACACCGGCGTCTGAGAATTATCATACTTGAACTCACTGGTC | Cloning TMD-truncated *Tom20-4* CDS into pBI221 vector carrying *nY* in the N terminus |

**Table S1 (continued). PCR primers for vector construction.**

| Primer name | Primer sequence (5′ to 3′) | Application |
| --- | --- | --- |
| Tom20-4-ΔTMD-2R | TTATGCGGCCGCTGCAGGTTACTGCCTTGACACCGGCGTCTGAGAATTATCATACTTGAACTCACTGGTC | Cloning TMD-truncated *Tom20-4* CDS into pGBKT7 vector |
| Tom20-4-TMD-F | GATCCGTGTTCGGATGGGTCATCTTAGCCAGTTACGTTGTTGCGTGGATCAGTTTTGCCTAAGGTAC | Cloning TMD of *Tom20-4* into in pBI221 vector carrying *nY* in the N terminus |
| Tom20-4-TMD-R | CTTAGGCAAAACTGATCCACGCAACAACGTAACTGGCTAAGATGACCCATCCGAACACG | |
| SP1-CDS-1F | CCAGGCCTACTAGTGGATCCATGATTCCTTGGGGTGGAGTTAC | Cloning *SP1* CDS into pBI221 vector carrying *nY* in the N terminus |
| SP1-CDS-1R | ACCCGGGAGCGGTACCTCAGTGACGATATGTCTTAACCGCC | |
| SP1-CDS-2F | TTGCTCCGTGGATCCGGTACCATGATTCCTTGGGGTGGAGTT | Cloning *SP1* CDS into HBT vector carrying *FLAG* tag in the C terminus |
| SP1-CDS-2R | CTTGTAGTCAGAAGGCCTCCCGGGGTGACGATATGTCTTAACCGC | |
| SP2-CDS-1F | CCAGGCCTACTAGTGGATCCATGGGAGCTCAGAAGAGTA | Cloning *SP2* CDS into pBI221 vector carrying *nY* in the N terminus |
| SP2-CDS-1R | ACCCGGGAGCGGTACCCTATGTTGATGAAGCAAGATTGGTG | |

**Table S1 (continued). PCR primers for vector construction.**

| Primer name | Primer sequence (5′ to 3′) | Application |
|---|---|---|
| RPN6-CDS-1F | CCAGGCCTACTAGTGGATCCATGGTTTCCTATCGTGCTACCACAG | Cloning *RPN6* CDS into pBI221 vector carrying *nY* in the N terminus |
| RPN6-CDS-1R | ACCCGGGAGCGGTACCTCAGGACATGATTTTGGCAGACCGG | |
| RPN6-CDS-2F | TTGCTCCGTGGATCCGGTACCATGGTTTCCTATCGTGCTACCACAG | Cloning *RPN6* CDS into HBT vector carrying *FLAG* tag in the C terminus |
| RPN6-CDS-2R | CTTGTAGTCAGAAGGCCTCCCGGGGGACATGATTTTGGCAGACCG | |
| HA-RPN6-CDS-F | AGAACACGGGGGACTCTAGAATGTACCCATACGATGTTCCAGATTACGCTATGGTTTCCTATCGTGCTACCACAG | Cloning *RPN6* CDS fused to *HA* tag in N terminus into pBI221 vector |
| RPN10-CDS-1F | CCAGGCCTACTAGTGGATCCATGGTTCTCGAGGCGACTATG | Cloning *RPN10* CDS into pBI221 vector carrying *nY* in the N terminus |
| RPN10-CDS-1R | ACCCGGGAGCGGTACCTCACTTCTTCTCATCCTCGCC | |
| RPN10-CDS-2F | TTGCTCCGTGGATCCGGTACCATGGTTCTCGAGGCGACTATG | Cloning *RPN10* CDS into HBT vector carrying *FLAG* tag in the C terminus |
| RPN10-CDS-2R | CTTGTAGTCAGAAGGCCTCCCGGGCTTCTTCTCATCCTCGCC | |
| HA-RPN10-CDS-F | AGAACACGGGGGACTCTAGAATGTACCCATACGATGTTCCAGATTACGCTATGGTTCTCGAGGCGACTATG | Cloning *RPN10* CDS fused to *HA* tag in N terminus into pBI221 vector |
| RPN12-CDS-1F | CCAGGCCTACTAGTGGATCCATGGATCCGCAGCTAACTGAAG | Cloning *RPN12* CDS into pBI221 vector carrying *nY* in the N terminus |
| RPN12-CDS-1R | ACCCGGGAGCGGTACCTTACACGATACGCTCCAGCTCTC | |
| RPN12-CDS-2F | TTGCTCCGTGGATCCGGTACCATGGATCCGCAGCTAACTGAAG | Cloning *RPN12* CDS into HBT vector carrying *FLAG* tag in the C terminus |
| RPN12-CDS-2R | CTTGTAGTCAGAAGGCCTCCCGGGCACGATACGCTCCAGCTCT | |
| RPN13-CDS-1F | CCAGGCCTACTAGTGGATCCATGAGTTCAAGCGAAGCGTTTC | Cloning *RPN13* CDS into pBI221 vector carrying *nY* in the N terminus |
| RPN13-CDS-1R | ACCCGGGAGCGGTACCTTAACTCTCATCCATCGCATCCC | |
| RPN13-CDS-2F | TTGCTCCGTGGATCCGGTACCATGAGTTCAAGCGAAGCGTTTC | Cloning *RPN13* CDS into HBT vector carrying *FLAG* tag in the C terminus |
| RPN13-CDS-2R | CTTGTAGTCAGAAGGCCTCCCGGGACTCTCATCCATCGCATCCCTTG | |
| CDC48A-CDS-1F | CCAGGCCTACTAGTGGATCCATGTCTACCCCAGCTGAATCTT | Cloning *CDC48A* CDS into pBI221 vector carrying *nY* or *cY* in the N terminus |
| CDC48A-CDS-1R | ACCCGGGAGCGGTACCCTAATTGTAGAGATCATCATCGTCCCC | |
| 6×His-CDC48A-CDS-F | AGAACACGGGGGACTCTAGAATGCATCATCATCATCATCATATGTCTACCCCAGCTGAATCTT | Cloning *CDC48A* CDS fused with *6×His* tag in N terminus into pBI221 vector |
| TTOPsgRNA-F | ATTGCTCTAGTAGCACCAATGCGT | Cloning sgRNA of *TTOP* into pKI1.1R vector |
| TTOPsgRNA-R | AAACACGCATTGGTGCTACTAGAG | |

**Table S1 (continued). PCR primers for vector construction.**

| Primer name | Primer sequence (5′ to 3′) | Application |
|---|---|---|
| TTOP-pGEX-F | <u>TCTGTTCCAGGGGCCCCTGG</u>GATCCATGGAAGATCAACCCATTAA | Cloning *TTOP* CDS into pGEX-GST vector |
| TTOP-pGEX-R | <u>TCAGTCAGTCACGATGC</u>GGCCGCTTATAGACCTTGATCATCTTTTAG | |
| Toc33NC-pET-F | <u>TGGATCGCATATGGGGC</u>CCGGGATGGGGTCTCTCGTTCG | Cloning C-terminally truncated *Toc33* CDS into pET507a-His vector |
| Toc33NC- pET-R | <u>GGATCTGATCAGGTACCC</u>TCGAGTTAGTCTACATGAATTGCTTTCCTC | |
| Toc33FL-pET-F | <u>TGGATCGCATATGGGGC</u>CCGGGATGGGGTCTCTCGTTCG | Cloning full-length *Toc33* CDS into pET507a-His vector |
| Toc33FL-pET-R | <u>GGATCTGATCAGGTACCC</u>TCGAGTTAAAGTGGCTTTCCACTTG | |
| Tom20-3NC-pET-F | <u>TGGATCGCATATGGGGC</u>CCGGGATGGATACGGAAACTGAGTT | Cloning the C-terminally truncated *Tom20-3* CDS into pET507a-His vector |
| Tom20-3NC-pET-R | <u>GGATCTGATCAGGTACCC</u>TCGAGTTAAGCATCATACTTGGCAT | |

The nucleotides underlined do not correspond to the target gene, but to the linker sequences.

**Table S. qRT-PCR primers.**

| Primer name | Primer sequence (5′ to 3′) |
|---|---|
| Toc33-F | TTGGATGTGTATAGAGTCGATGAGC |
| Toc33-R | CGTAGGAAAGTTCATCGGGAGGGGA |
| Toc34-F | ATCCAGCAGTTCCCACCTGCTACTC |
| Toc34-R | CTTTCCAACACCACCTTTCCCCATT |
| Toc159-F | ATCCCCGCCTCTCCCTTACTTGTTG |
| Toc159-R | GCTGGTCATACTCATCATCTTCCCC |
| Toc75-F | AGTTGCGGGAAGTAGAACAAGGGGC |
| Toc75-R | GACAGAGTACGGACCACCGAGGACA |
| PAP2-F | GCCTATTGCCTCGACAGTTCC |
| PAP2-R | CACACCACATTCGCCACCAC |
| Tic40-F | CACCTTTTCCGTTTCCATTTCC |
| Tic40-R | GTCTCTACTTTTGTCGCTGTCAC |

**Table S2. Protein and nucleotide sequences of coding-optimized used in this study**

| Constructs | Protein sequence | Nucleotide sequence |
|---|---|---|
| AtUba1 (72-1080) | EIDEDLHSRQLAV | GAGATTGACGAAGATCTGCACAGCCGTCAACTGGCGGTT |
| | YGRETMRRLFASNVLIS | TACGGTCGTGAAACCATGCGTCGTCTGTTCGCGAGCAACGTG |
| | GMHGLGAEIAKNLILAG | CTGATCAGCGGCATGCATGGTCTGGGCGCGGAAATCGCGAAA |
| | VKSVTLHDERVVELWDL | AACCTGATTCTGGCGGGTGTGAAGAGCGTTACCCTGCACGAC |
| | SSNFVFSEDDVGKNRAD | GAGCGTGTGGTTGAACTGTGGGATCTGAGCAGCAACTTCGTG |
| | ASVQKLQDLNNAVVVSS | TTTAGCGAGGACGATGTTGGCAAAAACCGTGCGGACGCGAGC |
| | LTKSLNKEDLSGFQVVV | GTTCAGAAGCTGCAAGATCTGAACAACGCGGTGGTTGTGAGC |
| | FSDISMERAIEFDDYCH | AGCCTGACCAAAAGCCTGAACAAGGAAGACCTGAGCGGTTTC |
| | SHQPPIAFVKADVRGLF | CAGGTTGTGGTTTTTAGCGATATCAGCATGGAGCGTGCGATT |
| | GSVFCDFGPEFAVLDVD | GAGTTCGACGATTACTGCCACAGCCACCAACCGCCGATCGCG |
| | GEEPHTGIIASISNENQ | TTCGTGAAGGCGGACGTTCGTGGTCTGTTTGGCAGCGTGTTC |
| | AFISCVDDERLEFEDGD | TGCGATTTTGGTCCGGAGTTCGCGGTGCTGGACGTTGATGGC |
| | LVVFSEVEGMTELNDGK | GAGGAACCGCACACCGGCATCATTGCGAGCATCAGCAACGAG |
| | PRKIKSTRPYSFTLDED | AACCAGGCGTTTATTAGCTGCGTTGACGATGAACGTCTGGAG |
| | TTNYGTYVKGGIVTQVK | TTCGAAGACGGTGATCTGGTGGTTTTTAGCGAGGTGGAGGGT |
| | QPKLLNFKPLREALKDP | ATGACCGAGCTGAACGACGGCAAACCGCGTAAGATCAAAAGC |
| | GDFLFSDFSKFDRPPLL | ACCCGTCCGTATAGCTTCACCCTGGACGAAGATACCACCAAC |
| | HLAFQALDHFKAEAGRF | TACGGCACCTATGTTAAGGGTGGCATTGTGACCCAGGTTAAA |
| | PVAGSEEDAQKLISIAT | CAACCGAAGCTGCTGAACTTTAAGCCGCTGCGTGAGGCGCTG |
| | AINTGQGDLKVENVDQK | AAGGACCCGGGTGATTTCCTGTTTAGCGACTTCAGCAAATTT |
| | LLRHFSFGAKAVLNPMA | GATCGTCCGCCGCTGCTGCACCTGGCGTTCCAGGCGCTGGAC |
| | AMFGGIVGQEVVKACSG | CACTTTAAAGCGGAAGCGGGTCGTTTCCCGGTGGCGGGCAGC |
| | KFHPLFQFFYFDSVESL | GAGGAAGATGCGCAAAAGCTGATCAGCATTGCGACCGCGATC |
| | PSEPVDSSDFAPRNSRY | AACACCGGTCAGGGCGACCTGAAAGTGGAGAACGTTGATCAA |
| | DAQISVFGAKFQKKLED | AAGCTGCTGCGTCACTTCAGCTTTGGCGCGAAGGCGGTTCTG |
| | AKVFTVGSGALGCEFLK | AACCCGATGGCGGCGATGTTTGGTGGCATTGTGGGTCAGGAA |
| | NLALMGVSCGSQGKLTV | GTGGTTAAAGCGTGCAGCGGCAAGTTCCACCCGCTGTTTCAA |
| | TDDDIIEKSNLSRQFLF | TTCTTTTACTTCGACAGCGTTGAGAGCCTGCCGAGCGAACCG |
| | RDWNIGQAKSTVAASAA | GTGGACAGCAGCGATTTTGCGCCGCGTAACAGCCGTTATGAC |
| | AVINPRFNIEALQNRVG | GCGCAGATCAGCGTTTTCGGTGCGAAATTTCAAAAGAAACTG |
| | AETENVFDDAFWENLTV | GAGGATGCGAAGGTGTTCACCGTTGGTAGCGGCGCGCTGGGT |
| | VVNALDNVNARLYVDSR | TGCGAATTTCTGAAAAACCTGGCGCTGATGGGCGTTAGCTGC |
| | CLYFQKPLLESGTLGTK | GGTAGCCAGGGCAAACTGACCGTGACCGACGATGACATCATT |
| | CNTQSVIPHLTENYGAS | GAGAAGAGCAACCTGAGCCGTCAGTTCCTGTTTCGTGACTGG |
| | RDPPEKQAPMCTVHSFP | AACATCGGCCAAGCGAAGAGCACCGTGGCGGCGAGCGCGGCG |
| | HNIDHCLTWARSEFEGL | GCGGTTATCAACCCGCGTTTCAACATTGAGGCGCTGCAAAAC |
| | LEKTPAEVNAYLSSPVE | CGTGTGGGTGCGGAAACCGAAAACGTTTTCGATGACGCGTTT |
| | YTNSMMSAGDAQARDTL | TGGGAAAACCTGACCGTGGTTGTGAACGCGCTGGACAACGTG |
| | ERIVECLEKEKCETFQD | AACGCGCGTCTGTACGTTGATAGCCGTTGCCTGTATTTTCAG |
| | CLTWARLRFEDYFVNRV | AAACCGCTGCTGGAAAGCGGCACCCTGGGCACCAAGTGCAAC |
| | KQLIYTFPEDAATSTGA | ACCCAAAGCGTTATCCCGCACCTGACCGAGAACTACGGTGCG |
| | PFWSAPKRFPRPLQYSS | AGCCGTGACCCGCCGGAAAAACAGGCGCCGATGTGCACCGTG |
| | SDPSLLNFITATAILRA | CACAGCTTCCCGCACAACATTGATCACTGCCTGACCTGGGCG |
| | ETFGIPIPEWTKNPKEA | CGTAGCGAGTTTGAAGGCCTGCTGGAGAAGACCCCGGCGGAA |
| | AEAVDRVIVPDFEPRQD | GTTAACGCGTACCTGAGCAGCCCGGTGGAGTATACCAACAGC |
| | AKIVTDEKATTLTTASV | ATGATGAGCGCGGGTGATGCGCAGGCGCGTGATACCCTGGAG |
| | DDAAVIDDLIAKIDQCR | CGTATCGTGGAATGCCTGGAGAAGGAAAAATGCGAAACCTTT |
| | HNLSPDFRMKPIQFEKD | CAAGATTGTTTAACCTGGGCGCGTCTGCGTTTCGAAGATTAC |
| | DDTNYHMDVIAGLANMR | TTTGTGAACCGTGTTAAACAACTGATTTATACCTTTCCGGAA |
| | ARNYSIPEVDKLKAKFI | GATGCGGCGACCAGCACCGGTGCGCCGTTCTGGAGCGCGCCG |
| | AGRIIPAIATSTAMATG | AAGCGTTTTCCGCGTCCGCTGCAGTACAGCAGCAGCGACCCG |
| | LVCLELYKVLDGGHKVE | AGCCTGCTGAACTTCATCACCGCGACCGCGATTCTGCGTGCG |

| | | |
|---|---|---|
| | AYRNTFANLALPLFSMA | GAAACCTTTGGTATCCCGATTCCGGAATGGACCAAAAACCCG |
| | EPLPPKVVKHRDMAWTV | AAAGAGGCGGCGGAAGCGGTTGACCGTGTGATCGTTCCGGAT |
| | WDRWVLKGNPTLREVLQ | TTCGAGCCGCGTCAAGACGCGAAAATTGTGACCGATGAAAAA |
| | WLEDKGLSAYSISCGSC | GCGACCACCCTGACCACCGCGAGCGTGGATGATGCGGCGGTT |
| | LLFNSMFTRHKERMDKK | ATCGATGACCTGATCGCGAAAATTGACCAGTGCCGTCACAAC |
| | VVDLARDVAKVELPPYR | CTGAGCCCGGATTTCCGTATGAAACCGATTCAATTTGAGAAG |
| | NHLDVVVACEDEDDNDV | GATGACGATACCAACTACCACATGGACGTTATCGCGGGTCTG |
| | DIPLVSIYFR | GCGAACATGCGTGCGCGTAACTATAGCATTCCGGAAGTGGAT |
| | | AAGCTGAAAGCGAAGTTCATCGCGGGTCGTATCATTCCGGCG |
| | | ATTGCGACCAGCACCGCGATGGCGACCGGCCTGGTGTGCCTG |
| | | GAGCTGTACAAAGTTCTGGACGGTGGCCACAAGGTGGAAGCG |
| | | TATCGTAACACCTTCGCGAACCTGGCGCTGCCGCTGTTTAGC |
| | | ATGGCGGAGCCGCTGCCGCCGAAAGTTGTGAAGCACCGTGAC |
| | | ATGGCGTGGACCGTGTGGGATCGTTGGGTTCTGAAAGGTAAC |
| | | CCGACCCTGCGTGAGGTTCTGCAGTGGCTGGAAGACAAGGGT |
| | | CTGAGCGCGTACAGCATCAGCTGCGGCAGCTGCCTGCTGTTC |
| | | AACAGCATGTTTACCCGTCACAAAGAGCGTATGGATAAGAAA |
| | | GTTGTGGACCTGGCGCGTGATGTGGCGAAGGTTGAACTGCCG |
| | | CCGTACCGTAACCACCTGGACGTTGTGGTTGCGTGCGAGGAT |
| | | GAAGACGATAACGACGTGGATATCCCGCTGGTTAGCATTTAT |
| | | TTCCGT |
| AtUbc8 | MASKRILKELKDL | ATGGCGAGCAAGCGTATCCTGAAGGAGCTGAAAGACCTG |
| | QKDPPTSCIFAGPVAED | CAGAAAGATCCGCCGACCAGCTGCATTTTTGCGGGTCCGGTG |
| (1-149) | MFHWQATIMGPAESPYS | GCGGAGGACATGTTCCACTGGCAAGCGACCATTATGGGTCCG |
| | GGVFLVTIHFPPDYPFK | GCGGAAAGCCCGTACAGCGGTGGCGTGTTCCTGGTTACCATT |
| | PPKVAFRTKVFHPNINS | CACTTTCCGCCGGACTATCCGTTCAAGCCGCCGAAAGTGGCG |
| | NGSICLDILKEQWSPAL | TTCCGTACCAAGGTTTTTCACCCGAACATCAACAGCAACGGT |
| | TISKVLLSICSLLTDPN | AGCATCTGCCTGGATATTCTGAAGGAACAGTGGAGCCCGGCG |
| | PDDPLVPEIAHMYKTDR | CTGACCATCAGCAAAGTGCTGCTGAGCATTTGCAGCCTGCTG |
| | AKYEATARNWTQKYAMG | ACCGACCCGAACCCGGACGATCCGCTGGTTCCGGAGATTGCG |
| | | CACATGTACAAGACCGATCGTGCGAAATATGAAGCGACCGCG |
| | | CGTAACTGGACCCAAAAATACGCGATGGGC |
| AtUb | MQIFVKTLTGKTI | ATGCAGATTTTTGTGAAGACCCTGACCGGCAAGACCATT |
| | TLEVESSDTIDNVKAKI | ACCCTGGAAGTTGAAAGCAGCGACACCATTGATAACGTGAAA |
| (1-76) | QDKEGIPPDQQRLIFAG | GCGAAAATCCAGGACAAAGAGGGTATCCCGCCGGATCAGCAA |
| | KQLEDGRTLADYNIQKE | CGTCTGATCTTCGCGGGTAAACAACTGGAAGACGGCCGTACC |
| | STLHLVLRLRGG | CTGGCGGATTACAACATTCAAAAGGAGAGCACCCTGCATCTG |
| | | GTTCTGCGTCTGCGTGGCGGT |
| SP1flex | GRSSGRDAEVLKT | GGGAGGTCAAGTGGAAGAGATGCTGAAGTACTTAAGACGGTC |
| | VTRVNQLKELAQLLELD | ACCCGTGTAAATCAGTTAAAAGAGTTGGCACAGTTGCTGGAG |
| (The RING | SKILPFIVAVSGRVGSE | TTAGACAGCAAGATCCTGCCATTTATCGTGGCCGTGAGCGGC |
| | TPIKCEHSGIRGVIVEE | CGTGTCGGCTCTGAGACCCCGATTAAATGCGAGCACAGTGGT |
| motif in | TAEQHFLKHNETGSWVQ | ATTCGTGGCGTGATTGTTGAAGAGACTGCGGAACAGCATTTT |
| | DSALMLSMSKEVPWFLD | TTAAAGCACAACGAAACGGGTAGCTGGGTTCAGGATAGTGCG |
| SP1RING is | DGTSRVHVMGARGATGF | TTGATGCTGAGCATGAGCAAGGAAGTGCCTTGGTTCCTGGAC |
| | ALTVGSEVFEESGRSLV | GACGGCACGAGCCGTGTGCATGTTATGGGTGCTCGTGGTGCG |
| shown in | RGTLDYLQGLKMLGVKR | ACCGGTTTCGCCTTGACCGTGGGTTCCGAGGTTTTTGAAGAA |
| | IERVLPTGIPLTIVGEA | AGCGGCCGTAGCTTGGTTCGCGGCACCCTGGACTATCTGCAA |
| bold) | VKDDIGEFRIQKPDRGP | GGTCTGAAGATGCTGGGTGTTAAGCGCATCGAGCGCGTCTTG |
| | FYVSPKSLDQLISNLGK | CCGACCGGCATCCCGCTGACCATTGTCGGTGAAGCAGTCAAG |
| | WGGGGSGGGGSHVIDSV | GATGATATTGGTGAATTTCGTATCCAGAAGCCGGACCGCGGA |
| | LERRRR**RQLQKRVLDAA** | CCGTTCTACGTTTCCCCGAAATCCCTGGACCAGCTGATCAGC |
| | **AKRAELESEGSNGARES** | AACCTGGGTAAATGGGGTGGCGGCGGTTCCGGCGGTGGCGGG |
| | **ISDSTKKEDAVPDLCVI** | TCGCACGTGATCGACTCAGTTCTAGAGCGTCGCCGTC**GTAGA** |
| | **CLEQEYNAVFVPCGMCC** | **CAACTGCAAAAACGTGTGCTCGACGCTGCGGCTAAACGCGCA** |
| | **CTACSSHLTSCPLCRRR** | **GAACTGGAGTCTGAGGGCTCCAACGGCGCGAGAGAAAGCATT** |
| | **IDLAVKTYRH** | **TCAGATAGCACCAAAAAAGAGGATGCGGTTCCGGATCTGTGT** |
| | | **GTTATTTGCCTGGAGCAAGAATACAATGCAGTATTCGTGCCG** |

| | | |
|---|---|---|
| | | **TGCGGTATGTGCTGCTGCACCGCGTGTAGCAGCCACCTGACT TCTTGTCCGCTCTGCCGTCGCCGTATCGATCTGGCGGTGAAG ACCTATCGTCAT** |
| RPN13 pru（9-122) | VMQEIMLEFRAGKMSLQ GTRVVPDARKGLVRIAR GDEGLIHFQWLDRNQNT VEDDQIVFPDEALFEKV NQSSDRVYILKFNSDDR KLFFWMQEPRAEGDAEL CSSVNQYLNQPL | gtaatgcaagaaataatgctagagtttagggcgggtaaaatg agcctgcagggtacgcgtgtagttccggatgcgcgtaagggc ttggttcgtattgcgcgtggtgatgaaggccttatccacttt cagtggctggatcgcaaccaaaacaccgtggaagatgatcag attgtctttccggacgaggctctgtttgaaaaggtgaaccaa tcgtccgaccgcgtgtatatcctgaaattcaacagcgacgac cgcaagctgttcttctggatgcaggagccacgtgcagagggc gacgccgagctgtgcagctctgttaatcagtacttgaatcaa ccgctg |
| TTOP UBL （23-99） | TLELNIKTLDSRTYTFQ VNKNETVLLFKEKIASE TGVPVGQQRLIFRGRVL KDDHPLSEYHLENGHTL HLIVRQPAE | actctagaattgaatataaaaacattagattcccgcacctat acctttcaggtcaataagaacgaaactgtgttgctgttcaaa gaaaagatcgcgagcgagacgggtgttccggttggccaacaa cgtctgattttccgcggtcgtgtgctgaaagatgaccacccg ctgagcgagtaccatttggagaacggccacaccctgcatctt atcgtgcgtcagccggcagaa |

**Table S3. PCR primers used for genotype identification.**

| Primer name | Primer sequence (5′ to 3′) | Application |
|---|---|---|
| TTOPpro-128F | CTGGAAATGTGTGTGGTAAAGG | For cloning of cDNA of TTOP to identify the TTOP knock-out mutant |
| TTOPgeno-519R | GCTTCCTCTACGGTGTAAGACTCA | |
| LB-TTOP | ATTTTGCCGATTTCGGAAC | LB, LP, and RP are used for PCR to identify the T-DNA |
| SALK_128909-LP | AGCTGAAGTCCCGCTATCTTC | |
| SALK_128909-RP | TATCCTATGTGAATGCAGGGC | |
| SALK_151742-LP | GCTTGACTTGTTGCAGGAAAC | |
| SALK_151742-RP | TTGCCTAGGTTGTGAATGACC | |

### Supplementary Materials and Methods

*Construction of Plasmids*

Most plasmids used in this study were generated using the One Step Cloning Kit (Vazyme) according to the manufacturer's instructions. All primers are listed in Table S1. All Arabidopsis coding sequences (CDSs) were PCR amplified from Col-0 cDNA. To generate constructs for yeast two-hybrid assays, full-length or truncated CDSs were amplified and inserted into the pGADT7 and pGBKT7 vectors (Clontech, USA). The constructs used for transient expression in Arabidopsis protoplasts were generated by cloning the PCR-amplified CDSs into a version of the pBI221 plasmid that was modified to contain the coding sequence of either *green fluorescence protein* (*GFP*), *yellow fluorescence protein* (*YFP*), or *mCherry* under the control of the cauliflower mosaic virus (CaMV) *35S* promoter. To generate constructs for protoplast transfection for co-immunoprecipitation (co-IP), CDSs were amplified with specific primers harboring the sequence encoding the Myc, 6xHistidine (His), or Hemagglutinin (HA) epitope tags and then cloned into the pBI221 or HBT vector. For BiFC assays, the full-length or truncated CDSs were cloned into the pBI221 vector containing either the sequence encoding the N-terminal half or the C-terminal half of YFP (nY or cY) as described previously (Waadt et al. 2008).

To generate the pBI121-*pTTOP:GFP-TTOP* construct to drive *TTOP* expression under the control of its own promoter, 1,983 bp of sequence upstream of the *TTOP* translation start was amplified and used to replace the original *35S* promoter in the pBI121 plasmid. The *GFP-TTOP* fragment was amplified from the plasmid pBI221-*p35S:GFP-TTOP* and cloned into the pBI121 plasmid containing the *TTOP* promoter.

To generate the pTA7002-*GFP-TTOP* construct for inducible transcription of *GFP-TTOP* by dexamethasone (DEX) treatment (Park et al. 2012), the *GFP-TTOP* fragment was amplified from pBI221-*p35S:GFP-TTOP* and cloned into the plasmid pTA7002. The same *TTOP* promoter fragment used above was cloned into pBI121 in place of the *35S* promoter to generate the pBI221-*pTTOP:GUS* construct. To generate the construct for generation of *ttop* mutants by genome editing using the CRISPR/Cas9 (clustered regularly interspaced short palindromic repeats/CRISPR-associated 9) system, a single guide RNA (sgRNA) with the sequence CTCTAGTAGCACCAATGCGT was cloned into the pKI1.1R vector, harboring the *RIBOSOMAL PROTEIN S5 A* (*RPS5A*) promoter, the *OLE1–TagRFP* (*red fluorescent protein*) cassette, and the *Cas9* endonuclease gene, as described previously (Pan et al. 2016). To generate the *p35S:mCherry-Toc33* and *p35S:mCherry-Tom20-2* constructs for plant transformation, the *mCherry-Toc33/Tom20-2* fragments were individually amplified and cloned into the Gateway® pENTR vector (Invitrogen, USA). The gene cassettes were then transferred into the pEarleyGate100 vector using Gateway LR Clonase (Invitrogen, USA) (Earley et al. 2006).

To generate vectors for protein production in *Escherichia coli*, the codon-optimized coding sequence of *SP1flex* (Ling et al. 2012), *AtUba1* (72-1080), *AtUbc8*, *AtUb* were ordered from GenScript (http://www.genscript.com), as listed in Table S2, and cloned into the pGEX-6p-1 vector (for GST-tagged fusion proteins). SP1 is a transmembrane protein and its overexpression in *E. coli* is prone to insolubility. To remove the transmembrane domain and express a soluble and catalytically active enzyme, the intermembrane domain (*SP1ims*, a.a. 21-225 of SP1) and the cytoplasmic domain (*SP1cyt*, a.a. 244-343 of SP1) were connected with a flexible linker: -[Gly4-Ser]2-) to create SP1flex in a previous study (Ling et al. 2012). To generate vectors for the production of recombinant Toc33FL (a.a. 1–297), Toc33NC (a.a. 1–251), TTOP (a.a. 1–879), Tom20-3NC (a.a. 1–174) and GST-

SP1RING, which carry the RING motif of E3 ligase (a.a. 254-343 of SP1) (Ling et al. 2012), in *E. coli*, all CDSs were PCR amplified from Arabidopsis Col-0 cDNA and were cloned into the pGEX-6p-1 vector (for GST-tagged fusion proteins) or the pET507a vector (for His-tagged fusion proteins) as listed in Table S1.

*Protein Production and Purification*

The expression vectors pGEX-6p-1-GST-SP1flex, pGEX-6p-1AtUba1, pGEX-6p-1-AtUbc8, pGEX-6p-1-AtUb, pGEX-6p-1-GST-SP1RING, pGEX-6p-1-GST-TTOP, pGEX-6p-1-GST-UBL, pGEX-6p-1-GST-RPN13pru, pET507a-His-Toc33NC, pET507a-His-Toc33FL, and pET507a-His-Tom20-3NC were transformed into *E. coli* BL21 cells (Sangon Biotech) for production of recombinant protein, which was induced by the addition of 0.4 mM isopropyl β-D-1-thiogalactopyranoside (IPTG) when the cultures reached $OD_{600}$ of 0.8. After incubation at 16°C for 18 h in the presence of IPTG, the cells were harvested by centrifugation at 14,160*g* for 12 min at 16°C. The pellets of cells carrying pGEX-6p-1 vector derivatives were resuspended in lysis buffer consisting of 30 mM Tris buffer pH 7.6, 500 mM NaCl, and 5 mM dithiothreitol (DTT) and sonicated on ice (Qsonica Q500 sonicator, 300 W, 10 s/10 s, 30 min on ice). After centrifugation at 29,000*g* for 36 min, the supernatant was filtered (0.22-μm Millipore filter) and then incubated with GSTrap Sepharose (GE Healthcare) for 2 h at 4°C. After extensive washing with wash buffer (30 mM Tris buffer, pH 7.6, 150 mM NaCl, and 5 mM DTT), the bound GST-tagged proteins were incubated with GST-tagged PreScission protease (GE Healthcare) at 4°C overnight to cleave the GST tag, and the target proteins without the GST tag were then collected in the eluted fractions. The pellets of cells harboring pET507a derivatives were resuspended in nickel binding buffer (30 mM Tris buffer, pH 7.6, 500 mM NaCl, and 1 mM Tris(2-carboxyethyl)phosphine [TCEP]) and sonicated on ice (Qsonica Q500 sonicator, 300 W, 10 s/10 s, 30 min on ice). After centrifugation at 29,000*g* for 36 min at 4°C, the supernatants were filtered (0.22-μm Millipore filter) and individually loaded onto a 5-mL HisTrap Ni-Chelating HP column (GE Healthcare). After extensive washing of the HisTrap column with 40 mM imidazole in nickel binding buffer, the His-tag proteins were eluted with 300 mM imidazole in nickel binding buffer.

For purification of His-Toc33FL, the cell pellet was resuspended in denaturing nickel binding buffer (30 mM Tris buffer, pH 7.6, 500 mM NaCl, 1 mM TCEP, 1% (v/v) TritonX-100 and 8 M urea) and sonicated on ice. After centrifugation at 29,000*g* for 36 min at 4°C, the supernatant was filtered (0.22-μm Millipore filter) and loaded onto a 5-mL HisTrap Ni-Chelating HP column (GE Healthcare). After extensive washing of the HisTrap column with 60 mM imidazole in denaturing nickel binding buffer, the fusion protein was eluted with 300 mM imidazole in denaturing nickel binding buffer. The solubilized recombinant His-Toc33 protein was first dialyzed in 200-fold (v/v) volumes of dialysis buffer 1 (30 mM Tris buffer, pH 7.6, 500 mM NaCl, 1 mM TCEP, 0.1% (v/v) TritonX-100 and 2 M urea) and then in buffer without urea, each time for 18 h at 4°C. After removal of precipitated proteins by centrifugation at 29,000*g* for 36 min at 4°C, soluble His-Toc33 was further purified by gel filtration using a HiLoad 26/60 Superdex 75 column (GE Healthcare) pre-equilibrated with 30 mM Tris, pH 7.5, 150 mM NaCl, and 2 mM DTT.

*In vitro Ubiquitination and Pull-down Assays*

*In vitro* ubiquitination assays were performed in a 30-μL reaction mixture. For the SP1 auto-ubiquitination assay, 1 μg of recombinant GST, GST-SP1flex, GST-SP1RING, or SP1flex was incubated with 0.1 μg of AtUba1 E1, 0.3 μg of AtUbc8 E2, and 2.5 μg of AtUb in a 30-μL reaction containing 30 mM Tris-HCl, pH 7.5, 2 mM ATP, 5 mM $MgCl_2$, 2 mM DTT, and 0.5 U inorganic pyrophosphatase (Sigma). The mixture was incubated at 30°C for 2 h, and the reaction was stopped by adding an equal volume of 2× sodium dodecyl sulfate-polyacrylamide gel electrophoresis (SDS-PAGE) buffer. Immunoblotting analysis was conducted using anti-ubiquitin antibodies (FK2, MERCK). To test the ubiquitination of substrates by SP1flex, the substrates His-Toc33NC, His-Toc33FL, and His-Tom20-3NC were individually added to the above reactions, and the reaction mixtures were subjected to immunoblotting analysis with anti-ubiquitin antibodies (FK2, MERCK). To test whether unknown E2/E3 ubiquitin conjugating enzymes or ligases in plant extracts can ubiquitinate His-Tom20-3NC, 20 µg of plant extract (PE) from 7-day-old seedlings was added to the *in vitro* ubiquitination assay.

In the pull-down analysis of ubiquitinated substrates by TTOP, 120 µL of ubiquitination reaction products was prepared using the above ubiquitination reaction system. The ubiquitination reaction products were incubated with equimolar amounts of GST or GST-TTOP in the presence of GSTrap Sepharose (GE Healthcare) in binding buffer (30 mM Tris, pH 7.5, 150 mM NaCl, and 2 mM DTT) for 1 h at 4°C. After extensive washing with binding buffer, GST-bound proteins were eluted with 20 mM reduced glutathione in binding buffer and subjected to immunoblotting analysis with anti-ubiquitin antibodies (FK2, Merck).

*Isothermal titration calorimetry (ITC) analysis*

ITC experiments were carried out on a VP-ITC Microcal calorimeter (Malvern) at 25 °C. All proteins were dissolved in buffer containing 50 mM Tris-HCl pH 7.5, 100 mM NaCl, and 1 mM DTT. Each titration point typically is consisted of injecting 10 µl aliquots of the RPN13 pru domain, at a concentration of 500 µM into the solution of containing TTOP UBL domain at a concentration of 50 µM. A time interval of 150 or 180 s between two titration points was used to ensure the complete equilibrium of each titration reaction. The titration data were analyzed using the program Origin7.0 and fitted by a one-site binding model.

*Protein Extraction, Co-immunoprecipitation, and Immunoblots*

For extraction of proteins from seedlings and protoplasts, total proteins were extracted in ice-cooled protein extraction buffer, containing 25 mM Tris-HCl, pH 7.5, 150 mM NaCl, 1 mM EDTA, 1% (v/v) Triton X-100, and 1% (v/v) plant protease inhibitor cocktail (P9599, Sigma, Germany). After incubation in extraction buffer for 30 min on ice, the supernatants were collected by centrifugation at 16,000*g* at 4°C for 15 min. The protein concentration in the supernatants was determined using the Bradford Protein Assay Kit (5000201EDU, Bio-Rad, USA).

For co-IP analysis, the transfected protoplasts were incubated overnight to express the transgenes. Total protein was extracted as mentioned above, and 80 µL of the supernatants was kept as input. The remaining supernatants were diluted with an equal volume of buffer containing 25 mM Tris-HCl, pH 7.5, 150 mM NaCl, 1 mM EDTA, and 1% (v/v) plant protease inhibitor cocktail [P9599, Sigma, Germany]), mixed with 10 µL of GFP-Trap agarose (gta-20, Chromotek, Germany), and incubated on a rotary mixer overnight at 4°C. The GFP-Trap agarose was used according to the manufacturer's instructions. After six washes of the GFP-Trap agarose with washing buffer (25 mM Tris-HCl, pH 7.5, 150 mM NaCl, and 1 mM EDTA), bound proteins were eluted from the GFP-Trap agarose by boiling in 2× SDS-PAGE buffer (50 mM Tris-HCl, pH 6.8, 20% glycerol, 1% SDS, and 0.1 M DTT) for 5 min and analyzed by SDS-PAGE and immunoblotting with anti-GFP and anti-FLAG antibodies.

For immunoblotting, the protein extracts were denatured in 2× SDS-PAGE buffer, separated on

SDS-PAGE gels, and transferred onto nitrocellulose membrane (GE10600002, Amersham™, Merck, Germany). SDS-PAGE and immunoblotting were performed as described (Pan et al. 2016) with minor modifications. When necessary, the gels were stained with Coomassie Brilliant Blue G250 (6104-58-1, Sigma, Germany). For immunoblots, the membranes were probed with the following primary antibodies, which were used according to the manufacturer's instructions: anti-GFP (SAB43011338, Sigma, Germany), anti-mCherry (PA534974, Thermo Fisher, USA), anti-TTOP (residues 158–449; the antigen for the immunization of rabbits was bacterially-produced, GenScript China Inc.), anti-HA tag (A02040, Abbkine, China), anti-Myc tag (A02060, Abbkine, China), anti-FLAG tag (A02010, Abbkine, China), anti-His tag (A02050, Abbkine, China), anti-cFBPase (AS04043, Agrisera, Sweden), and anti-actin (A01050, Abbkine, China). To identify TOC and TIC proteins, the following antibodies were used according to the manufacturer's instructions: anti-atToc75-III (AS08-351, Agrisera, Sweden), anti-atToc159 (ASO7239, Agrisera, Sweden), anti-atToc34 (AS07238, Agrisera, Sweden), anti-atToc33 (AS07236, Agrisera, Sweden), and anti-atTic40 (AS10709, Agrisera, Sweden). The following secondary antibodies were used according to the manufacturer's instructions: anti-rabbit IgG conjugated to horseradish peroxidase (170-6515, Bio-Rad [Shanghai]), anti-mouse IgG conjugated to horseradish peroxidase (A25012, Abbkine, China), anti-guinea pig conjugated to peroxidase (A5545, Sigma, Germany), and anti-chicken conjugated to peroxidase (A9046, Sigma, Germany). Chemiluminescence was detected using the ECL Plus Western Blotting Detection Reagents (1705061, Bio-Rad [Shanghai]) and an Alliance Q9 chemiluminescence and spectral fluorescence imaging system (UVITEC, UK). Signal intensities were quantified using ImageJ software (version 64-bit Java 1.8.0_172, USA) (Schneider et al. 2012).

The first-dimension BN-PAGE was performed as described previously (Eubel et al. 2005) with minor modifications. Transfected protoplasts were cultured overnight to allow transgene expression and lysed in native protein extraction buffer (30 mM HEPES, pH 7.4, 150 mM potassium acetate, 10% (v/v) glycerol, 1% (v/v) digitonin and 1% (v/v) plant protease inhibitor cocktail [P9599, Sigma, Germany], added before use). After incubation in extraction buffer for 30 min on ice, the supernatants were collected by centrifugation at 16,000*g* at 4°C for 15 min. The solubilized proteins were separated on a native gradient gel and transferred onto a nitrocellulose membrane, in both cases using 0.5× Tris borate EDTA (TBE) buffer containing 0.5% (w/v) Tris base, 0.04% (w/v) EDTA, and 0.3% (w/v) boric acid; they were then immunoblotted with anti-His, anti-HA, anti-FLAG, and anti-Myc antibodies as described above.

*GUS Staining and Transmission Electron Microscopy (TEM)*

Plant tissues were submerged in 90% (v/v) acetone fixation solution and incubated at 4°C for 40 min. The fixation solution was discarded, and the plant tissues were washed three times in wash solution (100 mM sodium phosphate buffer, pH 7.0, 10 mM EDTA, 2 mM potassium ferrocyanide, and 2 mM potassium ferricyanide). For GUS staining, the plant tissues were submerged in standard X-GlcA solution (100 mM sodium phosphate buffer, pH 7.0, 10 mM EDTA, 2 mM potassium ferrocyanide, 2 mM potassium ferricyanide, and 0.5 mg/mL X-GlcA), vacuum infiltrated for 5 min, and then incubated at 37°C for 4–6 h to allow development of the blue color, as described previously (Jefferson et al. 1987). The images of the GUS-stained specimens were captured using a stereomicroscope (model SZX16, Olympus, Japan). A minimum of five different samples were analyzed for each treatment, and typical images are shown.

The general procedures for transmission electron microscopy, sample preparation, and ultrathin sectioning were performed as described previously (Gao et al. 2012; Wang et al. 2012). The tissues from at least 10 plants per genotype were used for the TEM. Measurements of chloroplast cross-sectional areas were estimated using ImageJ software (version 64-bit Java 1.8.0_172, USA). At least 30 different plastids/chloroplasts per genotype were analyzed. The plastids were classified into three developmental stages: I, plastids with large rudimentary prothylakoids and prolamellar bodies (PLBs); II, plastids with PLBs of reduced sized and partially developed thylakoids; and III, plastids in which all PLBs had transformed into thylakoids.

*Phylogenetic analysis*

Amino acid sequences were obtained by BLAST searches of the Phytozome 13 database (Goodstein et al. 2012) (Table S4). Sequences were then aligned by the MAFFT multiple sequence alignment program (Katoh and Standley 2013). Poor quality alignments were filtered out by Gblocks (Castresana 2000) and only conserved sequences were retained. Phylogeny was inferred using IQ-TREE (Nguyen et al. 2015), and maximum likelihood tree was built. Bootstrap percentages at the branch points were estimated from 1,000 bootstrap replications. Finally, the phylogenetic tree was drawn by iTOL (Letunic and Bork 2016) using the tree file obtained from IQ-TREE.

*AlphaFold Prediction*

Modelled the binding of the TTOP UBL with RPN6 using the AlphaFold2 machine learning algorithm (Jumper et al. 2021; Mirdita et al. 2021). Take TTOP-UBL and RPN6 as an example, in the Google_colab AlphaFold2_advanced (https://colab.research.google.com/github/sokrypton/ColabFold/blob/main/beta/AlphaFold2_advanced.ipynb#scrollTo=rowN0bVYLe9n webpage, we input the TTOP-UBL and RPN6 sequences in the Sequence column and run 24 recycles for refinement
(TLELNIKTLDSRTYTFQVNKNETVLLFKEKIASETGVPVGQQRLIFRGRVLKDDHPLSEYHL
ENGHTLHLIVRQPAE:MVSYRATTETISLALEANSSEAITILYQVLEDPSSSPEAIRIKEQAITN
LCDRLTEEKRGEDLRKLLTKLRPFFSLIPKAKTAKIVRGIIDAVAKIPGTTDLQITLCKEMVEW
TRAEKRTFLRQRVEARLAALLMENKEYVEALALLSTLVKEVRRLDDKLLLVDIDLLESKLH
FSLRNLPKAKAALTAARTAANAIYVPPAQQGTIDLQSGILHAEEKDYKTGYSYFFEAFESFN
ALGDPRAVFSLKYMLLCKIMVSQADDVAGIISSKAGLQYVGPDLDAMKAVADAHSKRSLK
LFENALRDYKAQLEDDPIVHRHLSSLYDTLLEQNLCRLIEPFSRVEIAHIAELIGLPLDHVEK
KLSQMILDKKFAGTLDQGAGCLIIFEDPKADAIYSATLETIANMGKVVDSLYVRSAKIMS).
AlphaFold2 generated 5 models which were ranked by pLDDT (redicted Local Distance Difference Test) and pTMscore (post-translational modifications score) (Mariani et al. 2013; Tsaban et al. 2022). Higher pLDDT indicated higher prediction quality. But high pLDDT did not necessarily indicate a correct complex prediction, the inter-complex PAE (predicted alignment error) was used to rank complexes. Low inter PAE between chains indicates a confident prediction. The predictions of the interaction between TTOP UBL and RPNs, and the interaction between the C-terminal of TTOP with the transmembrane domains of TOC/TOM all followed the above method.

*Real-time quantitative PCR analysis*

Total RNA was extracted using the total RNA isolation TRIzol™ Reagent (Thermo Fisher Scientific, USA) following the manufacturer's instructions from 8-day-old *pTA7002-GFP-TTOP* plants subjected to DEX or DMSO treatment for 1 (D1) or 2 days (D2), or untreated. 1 μg total RNA was used for first-strand cDNA synthesis with the HiScript III Enzyme Mix, RT Mix, and oligo(dT)$_{20}$VN (Vazyme, China). Transcript levels were determined by RT-qPCR with the SYBR green master mix (Applied Biosystems, USA) on the StepOnePlus Real-Time PCR System (Applied Biosystems, USA). RT-qPCR was performed in triplicates. The relative expression levels were calculated with the $2^{-\Delta\Delta Ct}$ method (Livak and Schmittgen 2001). Primers used for RT-qPCR are listed in Supplementary Table 1.


Castresana J (2000) Selection of conserved blocks from multiple alignments for their use in phylogenetic analysis. **Molecular Biology and Evolution** 17: 540-552

Earley KW, Haag JR, Pontes O, Opper K, Juehne T, Song K, Pikaard CS (2006) Gateway-compatible vectors for plant functional genomics and proteomics. **The Plant Journal** 45: 616-629

Eubel H, Braun HP, Millar AH (2005) Blue-native PAGE in plants: a tool in analysis of protein-protein interactions. **Plant methods** 1: 11

Gao C, Yu CK, Qu S, San MW, Li KY, Lo SW, Jiang L (2012) The Golgi-localized Arabidopsis endomembrane protein12 contains both endoplasmic reticulum export and Golgi retention signals at its C terminus. **The Plant cell** 24: 2086-2104

Goodstein DM, Shu SQ, Howson R, Neupane R, Hayes RD, Fazo J, Mitros T, Dirks W, Hellsten U, Putnam N, Rokhsar DS (2012) Phytozome: a comparative platform for green plant genomics. **Nucleic Acids Research** 40: D1178-D1186

Jefferson RA, Kavanagh TA, Bevan MW (1987) GUS fusions: beta-glucuronidase as a sensitive and versatile gene fusion marker in higher plants. **The EMBO journal** 6: 3901-3907

Jumper J, Evans R, Pritzel A, Green T, Figurnov M, Ronneberger O, Tunyasuvunakool K, Bates R, Zidek A, Potapenko A, Bridgland A, Meyer C, Kohl SAA, Ballard AJ, Cowie A, Romera-Paredes B, Nikolov S, Jain R, Adler J, Back T, Petersen S, Reiman D, Clancy E, Zielinski M, Steinegger M, Pacholska M, Berghammer T, Bodenstein S, Silver D, Vinyals O, Senior AW, Kavukcuoglu K, Kohli P, Hassabis D (2021) Highly accurate protein structure prediction with AlphaFold. **Nature** 596: 583-589

Katoh K, Standley DM (2013) MAFFT Multiple Sequence Alignment Software Version 7: Improvements in Performance and Usability. **Molecular Biology and Evolution** 30: 772-780

Letunic I, Bork P (2016) Interactive tree of life (iTOL) v3: an online tool for the display and annotation of phylogenetic and other trees. **Nucleic Acids Research** 44: W242-W245

Ling Q, Huang W, Baldwin A, Jarvis P (2012) Chloroplast biogenesis is regulated by direct action of the ubiquitin-proteasome system. **Science** 338: 655-659

Livak KJ, Schmittgen TD (2001) Analysis of relative gene expression data using real-time quantitative PCR and the 2(-Delta Delta C(T)) Method. **Methods**: 402–408

Mariani V, Biasini M, Barbato A, Schwede T (2013) lDDT: a local superposition-free score for comparing protein structures and models using distance difference tests. **Bioinformatics (Oxford, England)** 29: 2722-2728

Mirdita M, Ovchinnikov S, Steinegger M (2021) ColabFold - Making protein folding accessible to all. **bioRxiv**: 2021.2008.2015.456425

Nguyen LT, Schmidt HA, von Haeseler A, Minh BQ (2015) IQ-TREE: A Fast and Effective Stochastic Algorithm for Estimating Maximum-Likelihood Phylogenies. **Molecular Biology and Evolution** 32: 268-274

Pan R, Satkovich J, Hu J (2016) E3 ubiquitin ligase SP1 regulates peroxisome biogenesis in Arabidopsis. **Proc Natl Acad Sci U S A** 113: E7307-E7316

Park C-J, Canlas PE, Ronald PC (2012) Establishment of glucocorticoid-mediated transcriptional induction of the rice XA21 pattern recognition receptor. **J Plant Biol** 55: 43-49

Schneider CA, Rasband WS, Eliceiri KW (2012) NIH Image to ImageJ: 25 years of image analysis. **Nature methods** 9: 671-675

Tsaban T, Varga JK, Avraham O, Ben-Aharon Z, Khramushin A, Schueler-Furman O (2022) Harnessing protein folding neural networks for peptide–protein docking. **Nature Communications** 13: 176

Waadt R, Schmidt LK, Lohse M, Hashimoto K, Bock R, Kudla J (2008) Multicolor bimolecular fluorescence complementation reveals simultaneous formation of alternative CBL/CIPK complexes in planta. **The Plant journal : for cell and molecular biology** 56: 505-516

Wang J, Tse YC, Hinz G, Robinson DG, Jiang L (2012) Storage globulins pass through the Golgi apparatus and multivesicular bodies in the absence of dense vesicle formation during early stages of cotyledon development in mung bean. **J Exp Bot** 63: 1367-1380
